# UniCell: Towards a Unified Solution for Cell Annotation, Nomenclature Harmonization, Atlas Construction in Single-Cell Transcriptomics

**DOI:** 10.1101/2025.05.06.652331

**Authors:** Luni Hu, Qianqian Chen, Ping Qiu, Hua Qin, Yilin Zhang, Lei Cao, Tianyi Xia, Ziqing Deng, Yong Zhang, Shuangsang Fang, Yuxiang Li

## Abstract

Standardizing cell type annotations across single-cell RNA-seq datasets remains a major challenge due to inconsistencies in nomenclature, variation in annotation granularity, and the presence of rare or previously unseen populations. We present UniCell, a hierarchical annotation framework that combines Cell Ontology structure with transcriptomic data for scalable, interpretable, and ontology-aware cell identity inference. UniCell leverages a multi-task architecture that jointly optimizes local and global classifiers, yielding coherent predictions across multiple levels of the ontology-defined hierarchy. When benchmarked across 20 human and mouse datasets, UniCell consistently outperformed state-of-the-art tools, including CellTypist, scANVI, OnClass, and SingleR, in annotation performance, and sensitivity to low-abundance populations. In disease settings, UniCell effectively identified previously unseen cell types through confidence-guided novelty detection. Applied to 45 human and 23 mouse tissue atlases, UniCell enabled cross-dataset and cross-species harmonization by embedding cells into a unified latent space aligned with Cell Ontology structure. Moreover, when used to supervise single-cell foundation models, UniCell substantially improved downstream annotation accuracy, rare cell detection, and hierarchical consistency. Together, these results establish UniCell as a generalizable framework that supports high-resolution annotation, nomenclature standardization, and atlas-level integration, providing a scalable and biologically grounded solution for single-cell transcriptomic analysis across diverse biological systems.

## Introduction

The rapid development of single-cell RNA sequencing (scRNA-seq) technology has provided unprecedented resolution for studying cellular heterogeneity, greatly accelerating our understanding of cell fate determination and functional specialization^1–3^. As a core step in single-cell analysis, cell type annotation forms the basis for interpreting cellular states and dynamic transitionss^4–6^. However, in practical applications, standardized annotation across datasets remains highly challenging due to technical variability in cell quality^7, 8^, the coexistence of shared and dataset-specific cell types^9, 10^, and inconsistencies in annotation granularity across laboratories^11, 12^. These issues significantly hinder the integrative analysis and reuse of single-cell data.

To address these challenges, international efforts such as the Human Cell Atlas (HCA))^13^, the Human BioMolecular Atlas Program (HuBMAP)^14^, and the Human Tumor Atlas Network (HTAN))^15^ have attempted to map cell type annotations to a unified terminology defined by the Cell Ontology (CO)^16^, thereby enabling the construction of reference atlases across tissues and studies. While these initiatives have promoted standardization of cell type nomenclature, most current approaches rely on manual alignment, which is inefficient, subjective, and difficult to scale. There is an urgent need for automated annotation frameworks that integrate transcriptomic data with structured prior knowledge, and support high-throughput, interpretable, and scalable analysis.

To address this need, a range of automated annotation tools have been developed, such as CellTypist^17^, which employs label transfer strategies; SingleR^18^, which relies on reference-based marker gene expression; and scANVI^19^, which utilizes deep generative modeling for probabilistic annotation. While these methods have improved annotation accuracy and efficiency, they generally adopt a flat classification strategy that fails to explicitly model hierarchical relationships among cell types^20–22^. As a result, their performance is limited when annotating complex cellular systems or identifying rare or previously unseen populations. For systems with clearly defined hierarchical structures, such as the immune system, conventional classifiers often fail to provide higher-level annotations when prediction confidence at finer granularity is low^23–28^.

Several studies have attempted to incorporate structural information into cell type annotation^29, 30^. For example, OnClass integrates the structural relationships defined in the Cell Ontology to expand the classification space and enhance rare cell type recognition^31^. However, such methods typically represent hierarchy using adjacency matrices and lack mechanisms for multi-level prediction, limiting their ability to capture hierarchical dependencies^32^. Other approaches rely on user-defined local trees, which are often not generalizable and heavily dependent on manual curation^33^.

Meanwhile, constructing cross-tissue and cross-species reference cell atlases is of great significance for understanding tissue specificity, developmental continuity, and evolutionary relationships among cell types^34–36^. With the rise of large-scale initiatives such as Tabula Sapiens^37^ and Tabula Muris^38^, the accumulation of multi-organ data has enabled the discovery of shared and tissue-specific cell types. However, existing annotation tools still suffer from low throughput, inconsistent naming conventions, and limited model generalizability, hindering the broader application of such reference atlases.

To overcome these limitations, we developed UniCell, a hierarchical cell type annotation framework guided by the Cell Ontology. UniCell explicitly models the hierarchical relationships among cell types and assigns each cell a standardized label along with its position within the ontology. This enables flexible annotation at varying levels of resolution and enhances the model’s capacity to identify ambiguous or previously unseen cell types. Applied across diverse human and mouse tissue datasets, UniCell was used to construct an atlas-level reference and demonstrated strong performance in nomenclature standardization, cross-dataset consistency, and novel cell type identification. To further extend its capability, we integrated UniCell with emerging single-cell foundation models (scFMs) such as scGPT^39^, GeneFormer^40^, and scFoundation^41^, which learn generalizable cell embeddings from large-scale single-cell data. By fine-tuning these embeddings under hierarchical supervision, UniCell achieves substantial improvements in rare cell detection, dataset generalization, and structured reasoning. This hybrid framework combining expert knowledge with foundation models offers a scalable, interpretable, and transferable solution for unified, ontology-aligned cell type annotation.

## Results

### UniCell: A Hierarchical Framework for Ontology-Guided Cell Identity Inference

We present UniCell, a hierarchical cell type annotation framework that integrates structured prior knowledge from the Cell Ontology with transcriptomic data to enable scalable, interpretable, and standardized cell identity inference (**Fig. 1a, top and Extended Data Fig. S1**). UniCell takes as input either raw or preprocessed gene expression matrices and optionally incorporates pretrained cell representations from single-cell foundation models (scFMs). These inputs are processed through a dedicated encoder to produce unified low-dimensional embeddings. The embeddings are subsequently propagated through a hierarchical classification module, in which each layer corresponds to a distinct level of the ontology-defined cell type hierarchy. Each level contains a fully connected neural network (FNN) that generates local probability distributions over candidate cell types via sigmoid activation. In parallel, global prediction heads output softmax-based terminal cell type probabilities and ordinal estimates of hierarchical depth. The entire model is trained using a multi-task learning objective that jointly optimizes local classification, global prediction, and hierarchy-aware supervision losses, ensuring consistent and biologically meaningful predictions across all levels of the ontology.

**Fig. 1.**
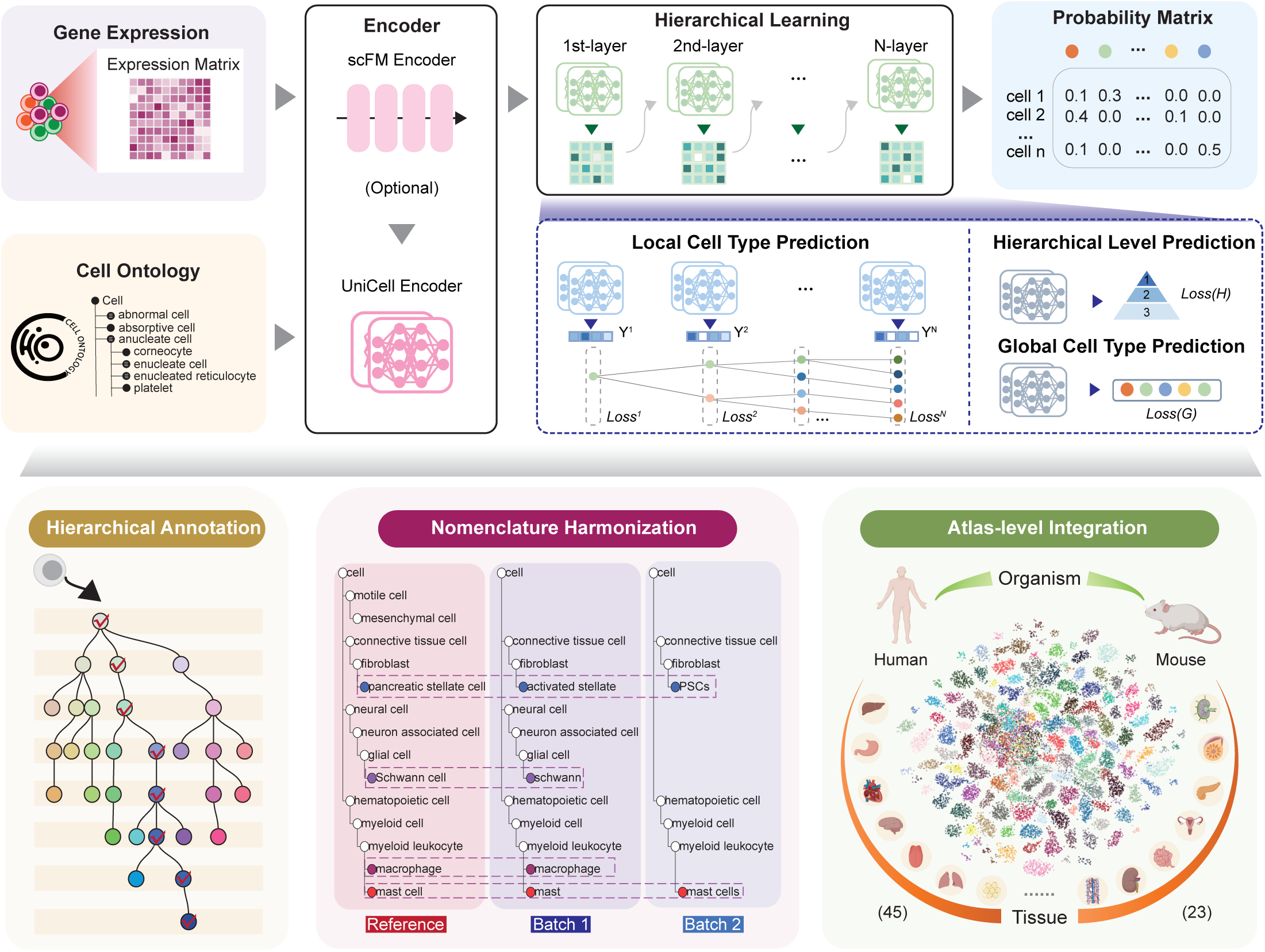
Schematic overview of UniCell. The framework integrates gene expression data and cell ontology for hierarchical cell type prediction and annotation. Top panel: Gene expression matrices are processed through an scFM encoder (optionally) followed by a UniCell encoder and then undergo hierarchical learning across multiple layers. A probability matrix is generated to represent cell type predictions. Local cell type predictions are refined through multiple hierarchical levels, optimizing loss functions at each stage (Loss¹, Loss², …, LossL). Hierarchical level and global cell type predictions further enhance classification accuracy. Bottom panel: The predicted cell types are applied to hierarchical annotation (left), nomenclature harmonization across datasets (middle), and atlas-level integration across species, enabling cross-organism comparisons (right).

UniCell’s architecture supports three integrated applications that address key limitations in large-scale single-cell analysis (**Fig. 1a, bottom**). First, for hierarchical annotation, UniCell explicitly models the Cell Ontology structure to assign each cell a path through the hierarchy, enabling both fine-grained and coarse-grained predictions. This layered annotation framework enhances interpretability and allows more informative outputs when the model encounters ambiguous or unseen cell types. Second, for nomenclature harmonization, UniCell projects cells from disparate datasets into a shared hierarchical embedding space, aligning inconsistent or dataset-specific labels to a standardized ontology. This enables cross-study label reconciliation and facilitates consistent annotation across batches, platforms, and studies. Third, for atlas-level integration, UniCell was applied to 45 human and 23 mouse tissue datasets to construct species-specific, mu lti-tissue reference atlases. These atlases demonstrate UniCell’s scalability and generalizability across biological systems, supporting high-resolution, ontology-aligned cell identity mapping across diverse single-cell datasets.

### UniCell Outperforms State-of-the-Art Methods in Cell Type Annotation

We evaluated UniCell in comparison with four commonly used annotation tools: CellTypist, OnClass, scANVI, and SingleR, across 20 representative single-cell RNA-seq datasets covering 10 human and 10 mouse tissues. UniCell consistently outperformed all methods across standard classification metrics, including accuracy, macro-F1 score, precision, and recall (**Fig. 2a**). Performance gains were especially pronounced in tissues with high cellular heterogeneity and complex lineage organization, such as human intestine, spleen, and mouse bone marrow (**Extended Data Fig. S2**). UniCell maintained robust annotation accuracy even under varying sequencing depths and differences in annotation granularity, underscoring its generalizability across diverse biological contexts.

**Fig. 2.**
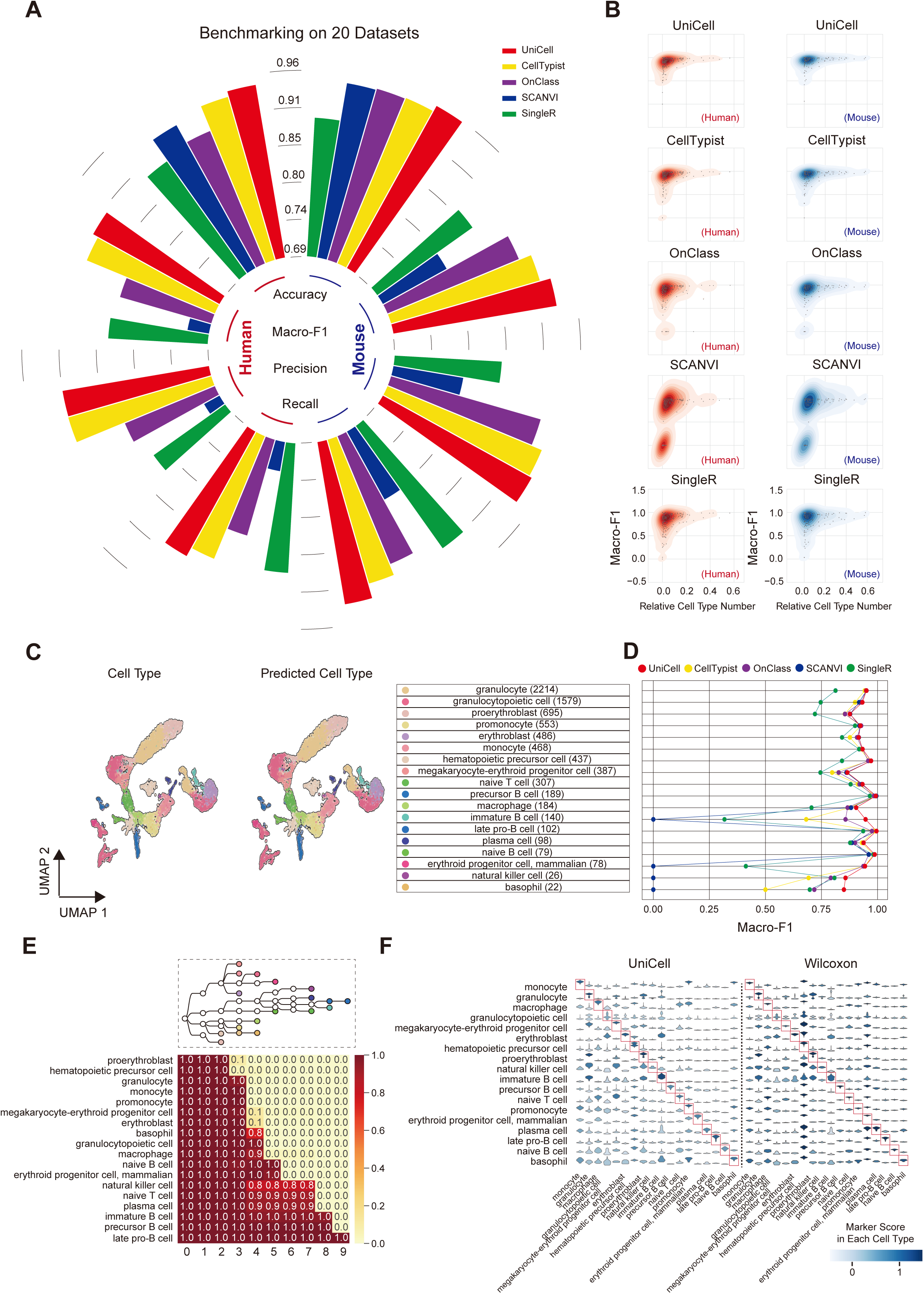
Evaluation of UniCell’s Annotation Performance. **a,** Radial bar chart showing the average annotation performance of UniCell, CellTypist, OnClass, SCANVI and SingleR across 10 human datasets and 10 mouse datasets. Metrics are divided into accuracy, Macro-F1, precision and recall. **b,** Density plot displaying the distribution of the Macro-F1 scores for annotation results from UniCell, CellTypist, OnClass, SCANVI, and SingleR across cell types with varying proportions of total cell numbers in the dataset. **c,** UMAP visualization of the ground truth labels (left) and predicted cell types in the mouse bone marrow dataset. **d,** Heatmap displaying the probability distribution of cell types across hierarchical levels in the mouse bone marrow dataset. The hierarchical tree diagram illustrates the hierarchical structure of cell types, constructed based on cell ontology. **e,** The top tree illustrating the hierarchical structure of cell types in the mouse bone marrow dataset, with the tree constructed based on cell ontology. The bottom heatmap displaying the probability of cell types across hierarchical levels. **f,** Stacked violin plots showing marker genes differentially expressed across cell types in the mouse bone marrow dataset. The left panel displays genes identified by the UniCell method, while the right panel shows those identified by the Wilcoxon method. Colors represent the marker score, calculated as the average expression of marker genes relative to the average expression of background genes.

UniCell also demonstrated superior sensitivity in annotating low-abundance cell populations. When performance was stratified by cell-type frequency, UniCell retained high macro-F1 scores for rare cell types (<100 cells), whereas competing methods exhibited substantial degradation in performance (**Fig.2b and Extended Data Fig. S3**). These findings indicate that UniCell’s hierarchical modeling is particularly advantageous for resolving sparsely represented populations, a critical challenge in tissue-level single-cell studies.

We next assessed the scalability and computational efficiency of UniCell. Across both human and mouse datasets, training time scaled linearly with dataset size (**Extended Data Fig. S4a,c**), and convergence in training loss, validation accuracy, and macro-F1 score was typically achieved within 5–10 epochs (**Extended Data Fig. S4b,d**). Even in large-scale datasets comprising over 90,000 cells, UniCell maintained per-epoch runtimes under one minute, enabling its practical application to high-throughput or atlas-level annotation pipelines. These results highlight the stability and tractability of the framework across a wide range of dataset complexities.

To evaluate UniCell’s capacity to capture biologically meaningful hierarchy, we analyzed its predictions in the mouse bone marrow dataset. UniCell accurately recovered terminal cell type labels in close agreement with expert annotations (**Fig. 2c**) and effectively identified rare types such as basophils and natural killer cells, which represent less than 1% of the total population, with macro-F1 scores exceeding 0.9 (**Fig. 2d**). Training dynamics further revealed rapid convergence for both common and rare populations, with macro-F1 scores stabilizing within the first 10 epochs (**Extended Data Fig. S5**). In addition to accurate label prediction, UniCell correctly mapped each cell type to its appropriate depth within the Cell Ontology hierarchy (**Fig. 2e**). Compared to existing approaches, UniCell embeddings more faithfully preserved known hematopoietic relationships, as reflected by increased intra-lineage similarity in the learned correlation structure (**Extended Data Fig. S6a**). Pseudotime analysis based on UniCell embeddings further revealed continuous developmental trajectories from hematopoietic precursors into distinct erythroid and lymphoid branches, recapitulating canonical hematopoietic hierarchies (**Extended Data Fig. S6b-d**). These results indicate that UniCell not only performs accurate classification, but also encodes interpretable and biologically coherent dynamic information.

Finally, we tested whether UniCell-derived annotations could enhance downstream biological inference. Using UniCell-predicted labels, we identified marker genes across multiple immune-related populations and performed gene ontology (GO) enrichment analysis. Compared to those defined using Wilcoxon-based annotations, UniCell-derived marker genes yielded a greater number of significantly enriched GO terms (**Fig. 2f, Extended Data Fig. S7a–b**), including lineage-relevant processes such as cytokine signaling, leukocyte activation, and hematopoietic differentiation. In particular, gene sets from monocytes, B-lineage cells, and hematopoietic progenitors demonstrated stronger enrichment for functionally consistent pathways (**Extended Data Fig. S7c**). Collectively, these findings show that UniCell provides not only accurate and hierarchical annotation but also biologically informative, interpretable gene-level outputs.

### UniCell Enables Identification of Previously Unseen Cell Types

Having systematically evaluated UniCell’s performance in conventional annotation tasks, we next investigated its ability to identify novel cell populations that emerge under pathological conditions. Such populations, often absent from training data, pose substantial challenges for reference-based annotation frameworks. To this end, we trained UniCell on healthy samples and evaluated its performance on disease-state cells. Across all disease contexts, UniCell consistently outperformed CellTypist, OnClass, scANVI, and SingleR, achieving the highest accuracy and macro-F1 scores (**Extended Data Fig. S8a**). In addition to label accuracy, UniCell embeddings preserved biological continuity between healthy and diseased states, enabling coherent representation of cellular hierarchies across conditions (**Extended Data Fig. S8b**).

In a representative case of follicular lymphoma, UniCell maintained superior annotation performance during the transition from normal lymph node to disease (**Extended Data Fig. S9a**). However, a distinct transcriptional subpopulation, originally annotated as follicular dendritic cells, was consistently misclassified by all methods, including UniCell, as a heterogeneous mixture of T cells, B cells, and plasma cells (**Extended Data Fig. S9b-c**). These cells exhibited high uncertainty in hierarchical predictions (**Extended Data Fig. S10a**) and inconsistent outputs from local classifiers (**Extended Data Fig. S10b-c**), illustrating the inherent limitations of reference-based approaches when encountering previously unseen types.

To address this challenge, we implemented a hierarchical confidence–based novelty detection pipeline. Specifically, query cells were embedded via PCA and clustered using the Leiden algorithm, followed by computation of within-cluster confidence based on UniCell-derived softmax probabilities. Clusters with low confidence and high internal consistency were flagged as candidate novel populations using Tukey’s fences (**Fig. 3a**). This approach identified Cluster 22 as a transcriptionally distinct outlier in the follicular lymphoma dataset (**Fig. 3b-d, Extended Data Fig. S11b**). Treating this cluster as a novel type significantly improved annotation performance across the dataset (**Fig. 3e**).

**Fig. 3.**
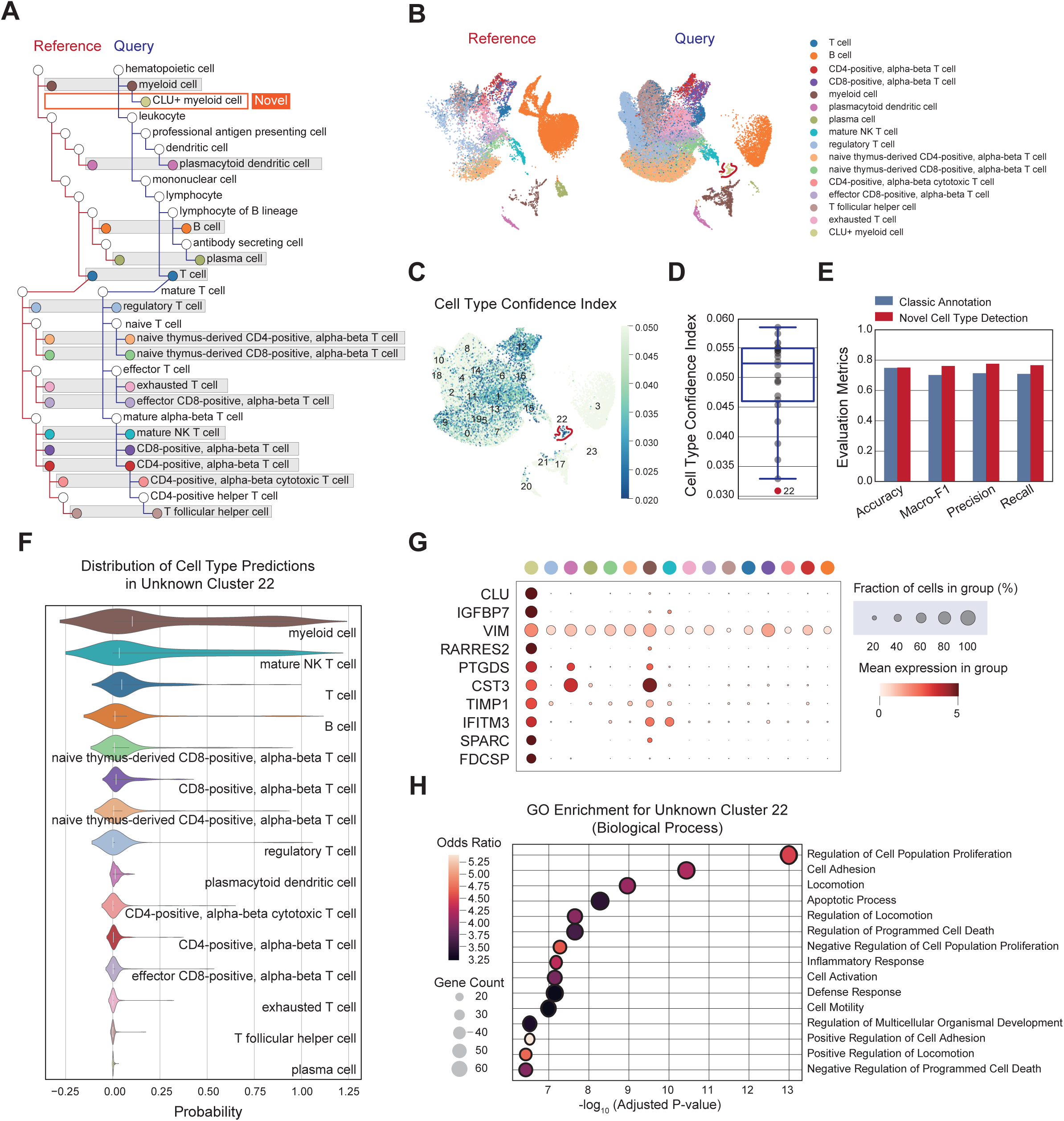
Identification and Characterization of Novel Cell Types Using Unicell. **a,** Hierarchical tree diagram representing the structure of cell types in the reference dataset, with orange branches highlighting novel cell types identified in the query dataset. **b,** UMAP visualization comparing the reference and query datasets, with colors indicating distinct cell types. **c,** UMAP visualization of the Cell Type Confidence Index across the dataset, where higher values represent more confident predictions of cell types. **d,** Bubble plot showing the expression levels of marker genes for Unknown Cluster 22, where dot size indicates the fraction of cells expressing each gene and color represents mean expression levels. **e,** Bar chart comparing evaluation metrics (accuracy, Macro-F1, precision, and recall) between classical annotation and novel cell type detection methods. **f,** Violin plot depicting the probability distribution of cell type predictions for Unknown Cluster 22, illustrating the uncertainty in classification. **g,** Bubble plot displaying significantly enriched biological processes from GO enrichment analysis for Unknown Cluster 22, with the size of the bubbles representing the odds ratio and color denoting adjusted p-values.

To characterize Cluster 22, we assessed its transcriptional similarity to reference populations. UniCell predictions indicated a closest match to myeloid cells (**Fig. 3f**), consistent with global correlation analysis between follicular dendritic cells and myeloid programs (**Extended Data Fig. S11c-d**). Marker gene identification using probability-weighted scores revealed highly specific expression of CLU, IGFBP7, VIM, and RARRES2 in Cluster 22 (**Fig. 3g, Extended Data Fig. S11e**). GO enrichment of these markers implicated pathways related to proliferation, adhesion, and immune activation (**Fig. 3h**), consistent with known follicular dendritic cell functions in tumor microenvironments.

Together, these results demonstrate that UniCell’s integration of hierarchical modeling and uncertainty estimation enables the detection, localization, and interpretation of previously unseen cell types, expanding the utility of cell annotation frameworks in disease settings.

### UniCell Performs Cross-Dataset Harmonization of Cell Type Labels

To further assess UniCell’s ability to harmonize cell type annotations across heterogeneous datasets, we applied it to human pancreas (hPancreas) and peripheral blood mononuclear cell (hPBMC) datasets. Leveraging a unified ontology-based reference, UniCell was trained across standardized annotated pancreas datasets and successfully aligned heterogeneous cell type labels to consistent hierarchical ontology nodes with high accuracy (**Fig. 4a-b**). To evaluate robustness across experimental protocols, we additionally applied UniCell to PBMC datasets generated using 10x Genomics 3′ and 5′ assay. Despite substantial technical variation, UniCell reliably matched biologically equivalent cell types across platforms and mapped them to unified ontology categories **(Extended Data Fig. S12a)**. Prediction confidence distributions further indicated that UniCell produced stable, well-calibrated assignments across multiple levels of the ontology hierarchy (**Extended Data Fig. S12b**).

**Fig. 4.**
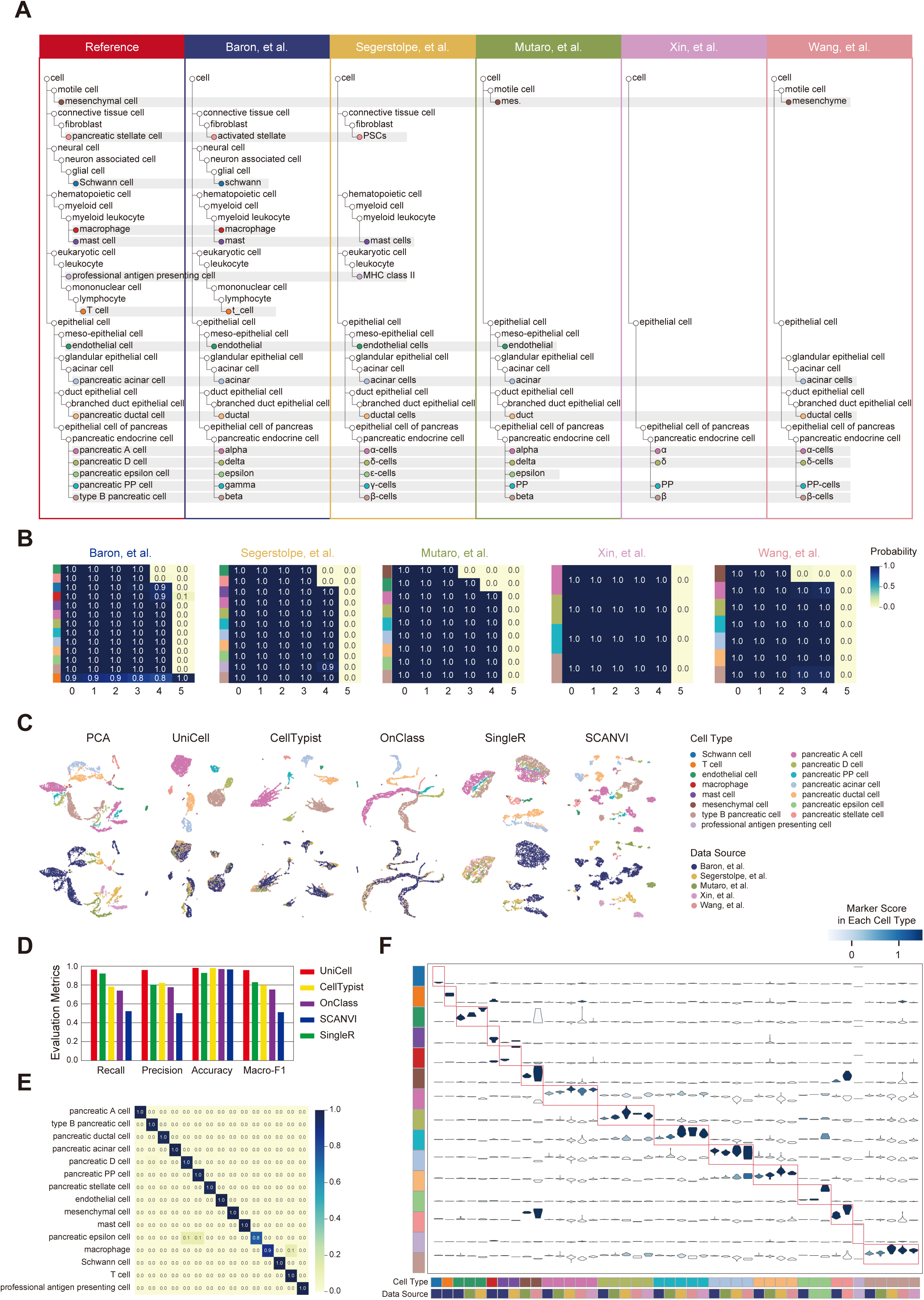
Harmonization of Cell Type Nomenclature Using UniCell. **a,** Hierarchical tree diagram representing the structure of cell types across different sources in the human pancreas (hPan) dataset, with differently colored regions highlighting cell types that have varying names across sources. **b,** Heatmap displaying the probability of cell types across hierarchical levels in different sources of the hPan dataset. **c,** UMAP visualizations comparing cell types (top) and data sources (bottom) in the hPan dataset, with embeddings derived from PCA, UniCell, CellTypist, OnClass, SingleR, and SCANVI. **d,** Bar chart comparing evaluation metrics (accuracy, Macro-F1, precision, and recall) for cell type annotation across UniCell, SingleR, CellTypist, OnClass, and SCANVI. **e,** Heatmap illustrating annotation accuracy across cell types in the hPan dataset using UniCell. **f,** Violin plot depicting the distribution of marker gene expression across cell types from various data sources in the hPan dataset, with red boxes highlighting distinct marker gene expression from each cell type across sources.

UniCell embeddings effectively preserved cell identity structure while mitigating batch effects. It achieved the highest average silhouette width (ASW) for cell type among all methods in both hPancreas and hPBMC datasets (**Extended Data Fig. S13a**). Correlation matrix analyses further confirmed that UniCell maximized intra-type similarity and minimized inter-type noise, yielding clearer block structures than SCANVI or SingleR **(Extended Data Fig. S13b-c**). These findings were consistent with UMAP projections, where UniCell embeddings showed improved cell-type separability and batch correction over PCA and competing tools (**Fig. 4c and Extended Data Fig. S13d**).

In addition to label harmonization, UniCell maintained high-resolution annotation capacity. Across pancreas and PBMC datasets, UniCell outperformed all other methods in accuracy, precision, recall, and macro-F1 (**Fig. 4d and Extended Data Fig. S14a**). Notably, it achieved 100% classification accuracy for dominant cell types, whereas other methods misclassified multiple subtypes (**Fig. 4e and Extended Data Fig. S14b**).

Finally, we assessed whether UniCell’s standardized labels improved downstream functional inference. Marker gene prediction showed higher specificity and interpretability across cell types in pancreas and PBMC datasets (**Extended Data Fig. S15a, c**). Moreover, UniCell successfully inferred markers for unannotated populations, such as canonical endocrine markers in pancreas and IL32 expression in mature αβ T cells from PBMCs (**Extended Data Fig. S15b, d**). These findings suggest that UniCell supports not only harmonized labeling but also robust and interpretable downstream functional analyses.

### UniCell Supports Atlas-Level Integration Across Species and Tissues

Building on UniCell’s demonstrated utility in capturing cell type features, , we next evaluated its capacity for large-scale cell atlas construction across species and tissues. We generated a comprehensive, ontology-based tree comprising over 400 human- and mouse-specific or shared cell types (**Fig. 5a**), and applied it as a unified reference for annotating millions of cells from both human and mouse atlases. UniCell consistently outperformed traditional annotation methods in accuracy and macro-F1 across both species (**Fig. 5b**), and maintained clear cell-type boundaries in low-dimensional embeddings (**Fig. 5c**), reflecting strong biological generalization across species.

**Fig. 5.**
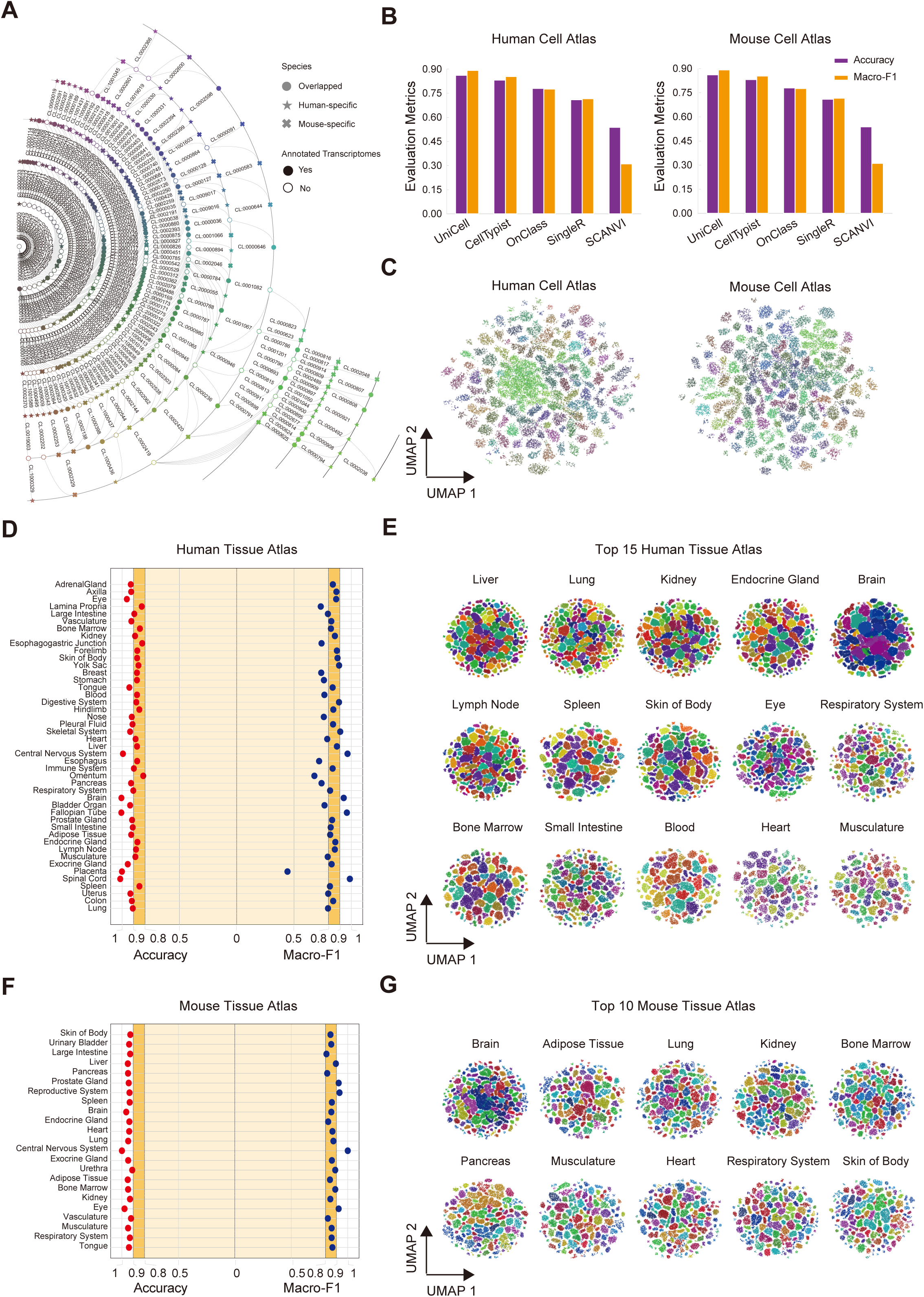
Construction of an Atlas-Level Reference Using UniCell. **a,** Hierarchical tree diagram representing the structure of cell types across human and mouse species, integrating data from multiple sources. **b,** Bar chart comparing the annotation performance (accuracy and Macro-F1) of UniCell, CellTypist, OnClass, SingleR, and SCANVI on human datasets. **c,** Bar chart comparing the annotation performance (accuracy and Macro-F1) of UniCell, CellTypist, OnClass, SingleR, and SCANVI on mouse datasets. **d,** Dumbbell plot showing accuracy and Macro-F1 scores for annotation results across 45 human tissue datasets. **e,** UMAP visualization of cell types from the top 15 human tissue datasets with the highest cell numbers used for UniCell training. **f,** Dumbbell plot showing accuracy and Macro-F1 scores for annotation results across 23 mouse tissue datasets. **g,** UMAP visualization of cell types from the top 10 mouse tissue datasets with the highest cell numbers used for UniCell training.

To further assess performance across tissues, we constructed tissue-specific ontologies for 45 human and 23 mouse organs (**Extended Data Fig. S17a, S18a**) and used these to annotate single-cell datasets with varying size, diversity, and complexity (**Extended Data Fig. S16**). Despite wide variability in gene number and cellular composition, UniCell achieved >0.9 accuracy in all mouse tissues and >0.8 in most human tissues (**Fig. 5d,f**), consistently outperforming alternative methods in macro-F1. UMAP projections revealed accurate clustering of functionally similar cells within each tissue, demonstrating UniCell’s capacity to capture tissue-specific expression programs (**Fig. 5e,g and Extended Data Fig. S17b, S18b**).

Collectively, these results indicate that UniCell enables ontology-driven, interpretable, and scalable annotation across species and tissues, wproviding a unifying framework for building comprehensive human and cross-species cell atlases.

### UniCell Enhances scFM Performance via Hierarchical Supervision

Beyond its standalone performance, we examined whether UniCell can serve as a structured supervision signal to improve single-cell foundation models (scFMs). We selected three leading scFMs—scGPT, GeneFormer, and scFoundation—and trained each using multi-level labels derived from UniCell’s hierarchical annotations. All models exhibited notable performance gains when supervised by UniCell (**Fig. 6a**). At the tissue level, scGPT+UniCell achieved improved annotation accuracy across most human tissues (**Fig. 6b**).

**Fig. 6.**
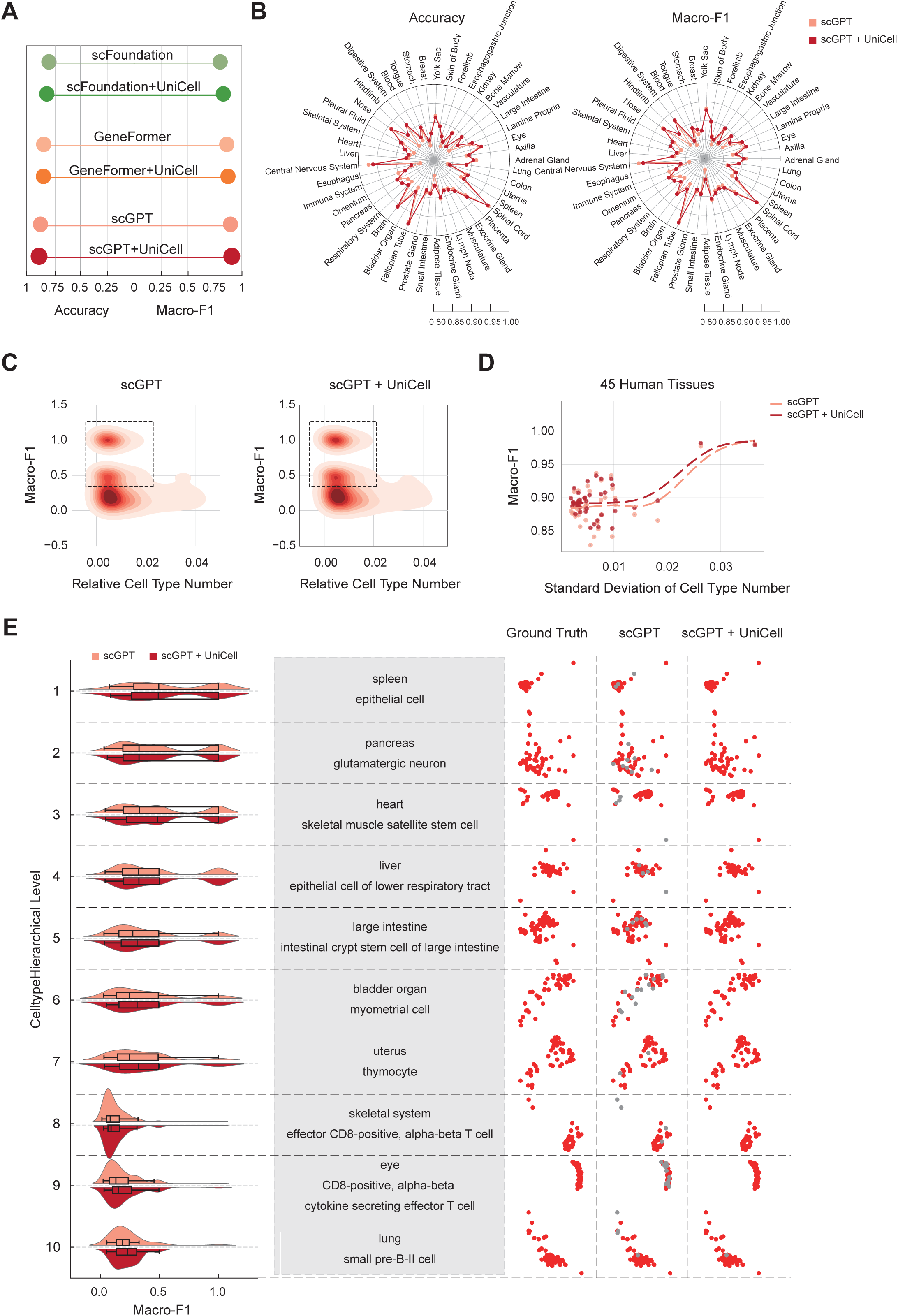
Integrating Single-Cell Foundation Models with UniCell. **a,** Dumbbell plot comparing the annotation performance (accuracy and Macro-F1) of large language models (scFoundation, GeneFormer, and scGPT) with their UniCell-enhanced counterparts. **b,** Radar chart comparing the annotation performance of scGPT with scGPT+UniCell across 45 human tissue datasets in terms of accuracy and Macro-F1. **c,** Density plot showing the distribution of Macro-F1 scores for scGPT and scGPT+UniCell annotation results across 45 human tissue datasets, stratified by cell type proportions. **d,** Line chart displaying the Macro-F1 score distribution for scGPT and scGPT_UniCell across 45 human tissue datasets, with standard deviations calculated based on cell type abundance. **e,** Violin plots showing the average Macro-F1 scores for scGPT and scGPT+UniCell across hierarchical cell type levels (left). A table listing selected cell types and their corresponding source tissues at each hierarchical level (middle). UMAP visualizations comparing ground truth labels with predictions from scGPT and scGPT+UniCell for the corresponding cell types in the table (right).

UniCell’s hierarchical supervision also improved model robustness against class imbalance. Compared to baseline scGPT, scGPT+UniCell showed higher macro-F1 scores for low-abundance cell types (**Fig. 6c**), and improved performance in tissues with high heterogeneity in cell-type proportions (**Fig. 6d**). In a stratified evaluation across ten representative cell types and ten ontology levels, the UniCell-supervised model outperformed the baseline at every level, with especially large gains for rare or structurally ambiguous types such as satellite muscle stem cells and uterine thymocytes (**Fig. 6e**).

We further assessed embedding quality using ASW scores, finding that all UniCell-supervised models achieved greater cell-type separation than their unsupervised counterparts (**Extended Data Fig. S19a,c,e**). UMAP projections showed better clustering of phenotypically similar cells (**Extended Data Fig. S19b,d,f**). At the tissue level, UniCell improved both ASW and classification metrics, particularly in highly heterogeneous contexts such as skin, muscle, and esophagus (**Extended Data Fig. S20a-b**). Across all ten ontology levels, scGPT+UniCell achieved widespread improvements in macro-F1 (**Extended Data Fig. S20c**), confirming the utility of hierarchical supervision for multi-resolution learning.

Together, these findings establish UniCell not only as a robust cell-type annotation tool but also as a powerful hierarchical supervisory signal for training scalable, generalizable, and biologically structured scFMs.

## Discussions

UniCell is a hierarchical, ontology-guided framework for cell type annotation that integrates structured prior knowledge with expression-based features within a multi-task learning architecture. This design enables strong generalization across datasets, species, and biological conditions. In contrast to widely used flat classification tools such as CellTypist, scANVI, and SingleR, UniCell outperforms across multiple evaluation metrics, particularly in rare cell identification, label harmonization, and novel population detection—substantially expanding the functional scope of automated annotation methods.

Conventional annotation frameworks typically rely on static reference labels or marker gene signatures, lacking the capacity to model the underlying hierarchical structure of cell types. As a result, they often fail in settings with ambiguous labels, heterogeneous resolution, or previously unseen populations. UniCell addresses these limitations by explicitly modeling the ontological hierarchy of cell types, enabling coarse-grained outputs when prediction confidence is low and providing biologically consistent, multi-resolution annotations via both global and local prediction heads. This structure-aware architecture not only improves classification accuracy but also facilitates the detection and interpretation of novel or transitional cellular states.

In terms of dataset integration, UniCell’s unified embedding space demonstrates robustness to batch effects, technical variation, and inconsistent annotation granularity. Its ontology-informed projections serve as a bridge between single-cell data and curated biological knowledge, offering a standardized framework for reference atlas construction. Nonetheless, while UniCell effectively reconstructs cross-organ and cross-species expression structures, its embedding stability under conditions of low gene expression or high technical noise remains to be systematically validated.

As a supervisory signal, UniCell’s hierarchical labels significantly enhance the performance of single-cell foundation models (scFMs) such as scGPT and GeneFormer. By providing structured supervision, UniCell improves model representation for rare, tissue-specific, and high-level cell types. This aligns with the emerging paradigm of “expert–foundation model synergy,” which seeks to combine domain-specific supervision with scalable pretrained architectures. The benefits of UniCell-guided supervision extend to downstream tasks such as cross-task transfer learning, cross-species generalization, and spatiotemporal trajectory inference.

Despite its strengths, UniCell has several limitations. Its reliance on the Cell Ontology implies that outdated or structurally incorrect entries may propagate errors to model outputs. Although the training process incorporates hierarchical depth supervision to mitigate this issue, more robust ontology modeling approaches, such as graph neural networks^42^ or learnable graph structures^43^, warrant further exploration. Additionally, UniCell currently operates as a static annotation framework and does not explicitly model temporal dynamics or developmental trajectories. Future extensions to dynamic graph-based modeling may enhance its applicability in developmental and regenerative systems.

In summary, UniCell offers a new paradigm for cell type annotation by unifying structure-aware inference, interpretable modeling, and scalable integration. Through continued integration with foundation models, biomedical ontologies, and temporal modeling frameworks, UniCell has the potential to serve as a core methodology for building multi-resolution, cross-species, and cross-condition reference cell atlases.

## Methods

### Unicell Model Design

#### Model Architecture

UniCell architecture consists of three core modules: Embedder, Hierarchical Learning, and Multi-task Learning, each contributing to cell feature extraction, hierarchical representation, and predictive learning.

#### Embedder

The embedding module is responsible for encoding single-cell gene expression data into a meaningful low-dimensional representation. Given an input expression matrix X_cell_ ∈ R^N×G^, where N is the number of cells and G is the number of genes, the UniCell encoder transforms the input into a latent feature space:

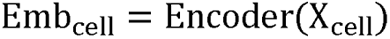

where Encoder() represents a neural network-based feature extractor, which could be a feedforward neural network (FNN), transformer-based model, or a pre-trained single-cell foundation model (scFM) that enhances feature extraction by leveraging prior biological knowledge. The resulting cell embeddings provide the foundation for subsequent hierarchical learning and classification tasks.

#### Hierarchical Learning

To effectively capture hierarchical relationships between cell types, the hierarchical learning module models the multi-level dependencies inherent in biological taxonomy. Rather than treating cell type annotation as a flat classification task, this module incrementally refines the feature representation across hierarchical levels.

For each hierarchical level l, the feature representation 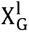 is determined by the output of the preceding layer 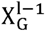 and the original cell feature Emb_cell_:

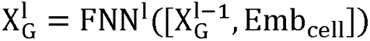

where FNN^l^ denotes the fully connected network at hierarchical level l, and 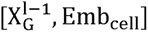 represents the concatenation of the previous layer’s output and the original cell embedding.

Moreover, ontology-based knowledge can be incorporated to inform hierarchical representation learning, thereby ensuring that the model aligns with established Cell Ontology by enforcing hierarchical consistency within the learned feature space.

#### Multi-task Learning

The multi-task learning module is designed to jointly optimize local cell type prediction and global hierarchical modeling:

##### Local Cell Type Prediction

At each hierarchical level, fully connected networks (FNNs) transform the feature representations and apply sigmoid activation for independent binary classification of cell types:

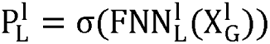

where FNN^l^ denotes the local classification network at level l, and σ represents the sigmoid function, which models the probability of a cell being assigned to a specific type at that hierarchical level.

##### Global Cell Type Prediction

The model synthesizes multi-level representations to learn a comprehensive global distribution of cell types, employing the softmax activation function for an all-encompassing classification of cell types:

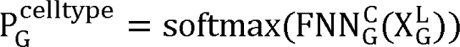

where 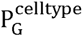 represents the probability distribution over all possible cell types, and L denotes the highest level in the hierarchy.

##### Global Hierarchical Prediction

A hierarchical level prediction task is incorporated to enhance the model’s comprehension of hierarchical structures:

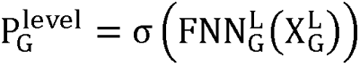

where 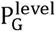 predicts the hierarchical level of a given cell, helping the model in capturing the hierarchical information of the cell.

##### Losses and Training

The UniCell model employs a weighted multi-task loss function to balance different hierarchical levels and learning objectives. The total loss consists of:

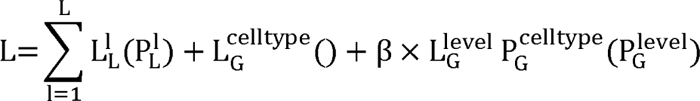

where 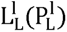 is the local classification loss at hierarchical level l, computed using binarycross-entropy (BCE) for multi-label classification; 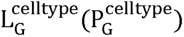 is the global classification loss, implemented as categorical cross-entropy for multi-class classification; 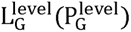 is the hierarchical level loss, enforcing consistency across hierarchical levels using BCE; β is a hyperparameter that controls the contribution of the hierarchical level loss.

### Data Processing

The Cell Ontology was first filtered to retain only is_a relationships, forming a directed acyclic graph. Node embeddings were then generated using Node2Vec (walk length = 3, dimension = 64, window size = 10, 100 epochs). Spearman correlations between embeddings were computed to assess semantic similarity among terms. For nodes with multiple parents, only the parent with the highest correlation was retained, and all other is_a edges were removed to simplify the ontology structure.

For all single-cell datasets, cells labeled as “unknown” or “cell” were removed. Data were normalized, log-transformed, and highly variable genes (HVGs) were selected (5,000 or 3,000 depending on the dataset). Unless otherwise noted, cells were randomly split into training, validation, and test sets in a 70:10:20 ratio. For normal-to-disease evaluation, “normal” samples were split 80:20 into training and validation sets, while “non-normal” samples were used for testing. In the Human Cell Atlas (HCA) and Mouse Cell Atlas (MCA), cells were stratified by annotated cell type and subsampled to reduce class imbalance: 100 cells (with replacement) for types with <100 cells; 500 cells (with replacement if <500) for types with 100–999 cells; and 1,000 cells for types with ≥1,000. For tissue atlases, all files from the same tissue were merged, retaining only primary data. To ensure balanced representation, up to 500 cells were sampled per cell type (with replacement if between 100–499 cells).

### Cell Type Annotation

UniCell performs supervised cell type annotation by training a classification model on labeled single-cell expression profiles. For each input cell, the model outputs a probability distribution over all candidate cell types via a softmax function, and the final prediction is assigned based on the class with the highest probability. Training was conducted using stratified data splits, and hyperparameters for each benchmarking task are listed in **Supplementary Table 1**. Annotation performance was evaluated across multiple human and mouse datasets using four standard metrics: accuracy, precision, recall, and macro macro-F1. To assess robustness across cell types, macro-F1 was computed per cell type and analyzed alongside cell abundance and relative representation, allowing evaluation across both major and rare populations.

### Novel Cell Type Detection

When applied to novel cell type detection, UniCell was trained on a reference dataset and tested on a separate query dataset containing potential unseen cell types. The detection procedure involved three steps. First, principal component analysis (PCA) was applied to the query data, and the top 20 components were used to construct a neighborhood graph, followed by Leiden clustering. Second, cell type probability distributions were computed using the softmax-transformed UniCell output, and the deviation of these probabilities was calculated within each Leiden cluster. Third, for each cluster, the median deviation and interquartile range (IQR) were used to identify low-variance clusters via Tukey’s fences. Clusters falling below the lower bound were flagged as outliers and considered candidate novel cell types.

### Nomenclature Harmonization

UniCell standardizes cell type annotations across datasets by mapping predictions to Cell Ontology terms, enabling consistent labeling across batches, platforms, and studies. To evaluate this capability, we applied UniCell to the human pancreas (hPancreas) and peripheral blood mononuclear cell (hPBMC) datasets, which comprise five and two batches, respectively. UniCell’s performance was benchmarked against CellTypist, OnClass, scANVI, and SingleR. Annotation accuracy was examined via annotation benchmarking metrics confusion matrices between predicted and ground truth labels, focusing on cell types shared across batches. We further assessed UniCell’s ability to preserve hierarchical relationships by comparing its predicted hierarchical distributions with the canonical structure in the Cell Ontology.

To quantify harmonization effectiveness in the embedding space, we computed Average Silhouette Width (ASW) scores for both cell types and batches using the scBI variant. The ASW for a single observation i is defined as:

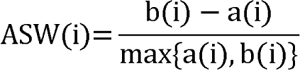

where a(i) is the average distance between i and all other points in the same cluster, and b(i) is the minimum average distance from i to points in other clusters. The final ASW score is the mean over all observations.

For cell types, ASW scores (ranging from 0 to 1) reflect the degree of cluster separation, with higher values indicating better-defined cell identities. For batches, ASW scores were inverted as ASW_batch_ =1 - ASW to reflect mixing quality, where 1 indicates ideal integration and 0 denotes complete batch separation. Lastly, we evaluated marker gene consistency across batches to determine whether UniCell could distinguish cell types in a biolo gically coherent and reproducible manner.

### Atlas-level Integration

To construct an atlas-level reference, we retrieved single-cell datasets from CellxGene and reclassified them by species and tissue type, resulting in 45 human and 23 mouse tissue-specific datasets (68 in total). Each dataset was processed and evaluated independently. UniCell models were trained separately on each dataset, and annotation performance was assessed using two standard classification metrics: accuracy and macro F1 score. To characterize dataset complexity and diversity, we recorded summary statistics for each dataset, including the number of source files merged, the number of annotated cell types, the total number of cells, and the number of genes retained after preprocessing.

### Single-cell Foundation Finetuning

To assess whether integrating UniCell with scFMs can enhance annotation performance, we incorporated pretrained embeddings from three leading scFMs—scGPT, GeneFormer, and scFoundation—into the UniCell framework (hereafter referred to as scFM+UniCell) and fine-tuned the model via full-parameter optimization on the cell type annotation task. Benchmarking was first performed on the HCA dataset, using the standalone versions of the scFMs as baselines.

To further investigate the contribution of scFM integration, we compared scGPT+UniCell with standalone scGPT across 45 human tissue datasets. Annotation performance was evaluated using accuracy and macro F1 score. Additionally, per–cell type performance metrics—including absolute counts and relative abundance—were collected, and the correlation between macro F1 scores and the standard deviation of cell counts was used to assess robustness to data imbalance.

To evaluate embedding quality, we extracted cell representations from both scFM+UniCell and standalone scFMs trained on HCA, as well as from scGPT+UniCell and scGPT trained on the 45 tissue datasets. Pairwise distance matrices were computed from these embeddings, and clustering quality was quantified using ASW scores, with ground truth cell type labels serving as reference.

Finally, to assess hierarchical consistency, we calculated macro F1 scores across multiple levels of the Cell Ontology for all 45 tissue datasets, comparing hierarchical annotation performance between scFM-integrated and standalone models.

### Comparison Approaches

To benchmark UniCell against existing cell type annotation methods, we compared its performance to four widely used classifiers: scANVI (v1.0.2, via scvi-tools), OnClass (v1.3), CellTypist (v1.6.2), and SingleR (v0.4.6). All methods were trained on the same datasets using their respective recommended workflows with default parameters (**Supplementary Table 4**). Training procedures followed publicly available pipelines: scANVI via scvi-tools, OnClass via its official documentation, CellTypist using the custom model training workflow, and SingleR using the standard pipeline.

We further evaluated the contribution of hierarchical supervision by comparing annotation performance using scFM embeddings with and without UniCell. In the baseline setting, scFM was fine-tuned on the training data using default settings, and its outputs were used directly for flat label prediction (**Supplementary Table 5**). In the integrated setting, scFM-derived embeddings were provided as input to UniCell, which applied multi-resolution supervision based on the Cell Ontology. Both configurations were trained on identical datasets and evaluated on the same test sets using accuracy, precision, recall, and macro F1 score, allowing assessment of performance gains attributable to hierarchical modeling.

### Inference of Cell Type–specific Marker Genes

To identify cell type–specific marker genes from UniCell outputs, we extracted the global classifier embeddings for each cell and transformed them into probabilistic cell type assignments. For numerical stability, the maximum value within each embedding vector was subtracted before applying the softmax function. Specifically, for a given cell i with raw logits z_i_, the probability vector over C cell types was computed as:

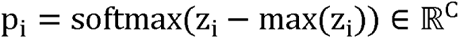

The raw single-cell RNA-seq matrix X ∈ ℝ^G×N^, containing expression values for G genes across N cells, was normalized, log-transformed, and capped at a maximum of 10. We then computed the cell type–specific gene embedding matrix M ∈ ℝ^G×C^ via a dot product between the transposed expression matrix and the stacked probability vectors across all cells:

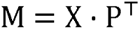

where P ∈ ℝ^C×N^ is the matrix formed by column-stacking the vectors p_i_ across all N cells. Each entry M_g,c_ quantifies the contribution of gene g to cell type c, weighted by cell-type assignment probabilities.

For nomenclature harmonization evaluation, we repeated this procedure using the unprocessed raw matrix X_raw_ in place of X, allowing marker inference in the native expression space:

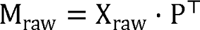

As a baseline comparison, we ranked genes using the Wilcoxon rank-sum test, identifying differentially expressed genes across cell types based on annotated training labels. The resulting rank scores served as an independent estimate of each gene’s discriminatory power.

### Trajectory Inference

To assess whether UniCell captures biologically meaningful cell hierarchies, we performed pseudotime trajectory inference using its cell embeddings in the mouse bone marrow dataset. Mean embeddings were computed for each annotated cell type, and pairwise Pearson correlations between these averages formed a similarity matrix, from which a maximum spanning tree was constructed to infer global cell type relationships. Independent component analysis (ICA) was applied to both the correlation matrix and the full UniCell embedding matrix to visualize lineage structure and infer pseudotemporal dynamics. Based on biological priors, five early-stage cell types, hematopoietic precursor cell, megakaryocyte–erythroid progenitor cell, erythroid progenitor cell (mammalian), granulocytopoietic cell, and proerythroblast, were used as source states. Mean embeddings from these sources were used to compute cosine distances to all cells; the minimum distance per cell defined its pseudotime, which was rescaled by setting the global minimum to zero. As a benchmark, the same procedure was performed using PCA-derived components to compare UniCell-based pseudotime inference with PCA in capturing lineage continuity.

### GO Enrichment

To assess the biological relevance of cell type–specific marker genes identified by UniCell, we performed Gene Ontology (GO) enrichment analysis using the gseapy.enrichr() function and the GO collections from MSigDB 2024.1. The top 100 marker genes per cell type were used as input, with B cell subtypes pooled into a single B lineage group in the mouse bone marrow dataset. As a benchmark, the same enrichment analysis was applied to marker genes identified using the Wilcoxon rank-sum test.

## Data availability

All data used in thes research were collected from published sources, with detailed descriptions provided in **Supplementary Table 1-2**.

## Code availability

UniCell is packaged, and distributed as an open-source, publicly available repository at https://github.com/BGIResearch/UniCell.

## Funding

This work is supported by the National Natural Science Foundation for Young Scholars of China (32300526) and National Key R&D Program of China (2022YFC3400400).

## Acknowledgement

We acknowledge the Stomics Cloud platform (https://cloud.stomics.tech/) for providing GPU computational resources. We thank the colleagues in our research group for inspiring discussion and their contributions.

## Author contributions

Y.L., S.F., Y.Z., and L.H conceptualized the study. L.H. and P.Q. were responsible for the model design and tool implementation. L.H. and Q.C. performed data analysis and model evaluation. H.Q., Y.L., L.C.,T.X. and Z.D. contributed key ideas and advice. L.H., Q.C. and P.Q. wrote the manuscript. Y.L., S.F., and Y.Z. supervised the study.

---

**Extended Data Fig. 1 | Hierarchical Cell Representation Learning Model with Multi-Task Learning.** The model comprises three main components: cell embeeding, cell hierarchical learning and multi-task learning. Gene expression data are first encoded into cell embeddings () via an encoder. An optional large language model (LLM) can be used to enhance the embedding representation. The embeddings are then processed through a series of feedforward neural networks (FNNs) in a hierarchical manner, generating intermediate representations (, , …,). In the multi-task learning framework, both local and global learning layers are incorporated. Local layers utilize FNNs to process hierarchical representations (, , …,) to predict local cell types (, , …,) using sigmoid activation. Global layers consist of two FNNs predict global cell properties: one employing softmax activation to classify global cell types (), and the other using sigmoid activation to infer hierarchical cell levels ().

**Extended Data Fig. 2 | Annotation Performance Across Multiple Datasets.** 3D surface plots illustrating the annotation performance of UniCell, CellTypist, OnClass, SCANVI and SingleR across 10 human datasets (**a**) and 10 mouse datasets (**b**). Metrics are divided into accuracy, Macro-F1, precision and recall.

**Extended Data Fig. 3 | Relationship Between Cell Type Abundance and Annotation Performance.** Line charts displaying the GAM-fitted mean Macro-F1 score against the number of cells per cell type across 10 human datasets (**a**) and 10 mouse datasets (**b**).

**Extended Data Fig. 4 | Training Efficiency and Performance Evolution. a,** Scatter plot showing the relationship between mean training runtime per epoch (s) and the number of cells across 10 human datasets. **b,** Line chart depicting the evolution of training loss (blue), annotation accuracy (green), and Macro-F1 score (red) across epochs for the 10 human datasets. Solid lines represent mean values, and shaded regions show metric distribution. **c,** Scatter plot showing the relationship between mean training runtime per epoch (s) and the number of cells across 10 mouse datasets. **d,** Line chart displaying the evolution of training loss, annotation accuracy, and Macro-F1 score across epochs for the 10 mouse datasets. Solid lines represent mean values, and shaded regions indicate metric distribution.

**Extended Data Fig. 5 | Evolution of Macro-F1 Scores Across Training Epochs. a,** Line charts displaying the evolution of Macro-F1 scores for specific cell types over 30 training epochs. Titles include the corresponding cell types, their cell numbers, and proportions in the dataset.

**Extended Data Fig. 6 | Analysis of Cell Type Relationships and Pseudotime. a,** Heatmaps showing the correlation of cell types using latent spaces derived from UniCell, PCA, CellTypist, SCANVI, OnClass, and SingleR. **b,** NNetwork visualizations depicting the ordered connections between cell types, as inferred by UniCell (top) and PCA (bottom). **c,** Scatter plots of the UniCell (top) and PCA (bottom) latent spaces, projected into two dimensions using independent component analysis (ICA), with cells colored by the cell type labels and pseudotime. **d,** Violin plots displaying the pseudotime distribution of cell types, as inferred by UniCell (top) and PCA (bottom).

**Extended Data Fig.7 | Differential expressed genes analysis and functional enrichment analysis of the mouse bone marrow dataset. a,** Bubble plots showing marker genes identified by the Unicell method (top) and the Wilcoxon test (bottom). Dot sizes represent the proportion of cells expressing each marker gene, while colors indicate expression levels. **b,** Bar plot comparing the number of enriched GO pathways identified by the Unicell method and the Wilcoxon test. **c,** Bubble plot displaying enrichment results across different cell types and GO pathways. Dot sizes represent the number of genes enriched in each pathway per cell type, and colors denote statistical significance.

**Extended Data Fig.8 | Annotation Performance from Normal to Disease Across Multiple Conditions. a,** Radial bar charts showing the annotation performance using normal cells as reference and disease cells as query across various disease conditions for UniCell, CellTypist, OnClass, SCANVI, and SingleR. The left panel illustrates Accuracy, and the right panel displays Macro-F1 scores across multiple diseases. **b,** UMAP visualizations of single-cell data from multiple disease conditions. Each row corresponds to a distinct dataset, with cells colored by predicted cell types (left column) and annotated disease states (right column).

**Extended Data Fig.9 | Cell Type Annotation Benchmarking in Follicular Lymphoma. a,** Bar chart comparing evaluation metrics (accuracy, Macro-F1, precision, and recall) for cell type annotation across UniCell, SingleR, CellTypist, OnClass, and SCANVI, when using normal condition as reference and follicular lymphoma condition as query. **b,** Heatmaps illustrating annotation accuracy across cell types in the follicular lymphoma condition using UniCell and other annotation methods. **c,** UMAP visualizations of the follicular lymphoma condition, colored by cell type. The top-left panel displays the ground truth annotations, while the remaining panels show predicted cell type labels from each model.

**Extended Data Fig.10 | Hierarchical and Local Cell Type Predictions in Follicular Lymphoma Condition. a,** Heatmap showing hierarchical prediction probabilities across hierarchical levels for each true cell type. **b,** Heatmap of local cell type prediction probabilities, with true cell types on the y-axis and predicted labels on the x-axis. **c,** Heatmap displaying local prediction probabilities for 108 follicular dendritic cells, which were not included in the reference.

**Extended Data Fig. 11 | Characterization of a Novel Cell Type Identified in Follicular Lymphoma. a,** UMAP visualizations of the follicular lymphoma condition, colored by Leiden clusters. **b,** Violin plot showing the distribution of cell type confidence index across Leiden clusters. **c,** Heatmap of pairwise Pearson correlation coefficients between cell types based on gene expression profiles**. d,** Heatmap of global cell type prediction probabilities. **e,** Volcano plot displaying differentially expressed genes in Cluster 22 compared to other clusters. Red points represent significantly upregulated genes (adjusted p < 0.05, |logLFC| > 1). **f,** Heatmap of relative expression of highly expressed genes in Cluster 22 across reference immune cell types.

**Extended Data Fig.12 | Hierarchical Alignment of Cell Types Across hPBMC Data Sources. a,** Panel showing the hierarchical structure of cell types across different sources of the human peripheral blood mononuclear cell (hPBMC) dataset. Dashed lines indicate correspondences between cell types with different names across sources. **b,** Heatmaps showing prediction probabilities for cell types across hierarchical levels, comparing annotations from different sources within the hPBMC dataset.

**Extended Data Fig.13 | Comparison of Latent Space Derived from Annotation Methods. a,** Scatter plots comparing Average Silhouette Width (ASW) scores for PCA, UniCell, CellTypist, OnClass, SingleR, and SCANVI on the hPancreas (left) and hPBMC (right) datasets. The x-axis represents ASW scores for data source separation, and the y-axis represents ASW scores for cell type separation. **b,c,** Heatmaps of pairwise Pearson correlation coefficients between cell types based on gene expression profiles in the hPancreas (**b**) and hPBMC (**c**) datasets, showing cell type similarity across methods. **d,** UMAP visualizations of the hPBMC dataset, colored by cell type (top) and data source (bottom), for embeddings generated by PCA, UniCell, CellTypist, OnClass, SingleR, and SCANVI.

**Extended Data Fig.14 | Annotation Performance on the hPBMC Dataset. a,** Bar plots comparing annotation performance across UniCell, SingleR, CellTypist, OnClass, and SCANVI on the hPBMC dataset. Evaluation metrics include Accuracy, Macro-F1, Precision, and Recall. **b,** Heatmap illustrating annotation accuracy per cell type for UniCell, highlighting its performance consistency across the hPBMC dataset.

**Extended Data Fig.15 | Marker-Inference in hPancreas and hPBMC Datasets. a,** Heatmap showing expression of selected marker genes across batches and cell types in the hPancreas dataset. **b,** Dot plot displaying inferred marker genes for pancreatic endocrine cell. Dot size indicates the fraction of cells expressing the gene, and color reflects mean expression levels. **c,** Dot plot summarizing marker gene expression across cell types in the hPBMC dataset. Dot size indicates the fraction of cells expressing the gene, and color reflects mean expression levels. **d,** UMAP visualization of the expression levels of inferred marker genes (IL32, MAL, CD8A) used to distinguish mature alpha-beta T cell, CD4-positive, alpha-beta T cell and CD8-positive, alpha-beta T cell.

**Extended Data Fig. 16 | Overview of Human and Mouse Datasets Used for Tissue Atlas Construction.** Bubble plots showing the overview of the human tissue datasets (**a**) and the mouse tissue datasets (**b**) for construction of the across-species atlas-level reference. Dot size represents the number of files comprising each dataset, while color indicates the number of cell types within each dataset.

**Extended Data Fig.17 | Hierarchical Structure and Visualization of Cell Types in the Human Tissue Atlas. a,** Hierarchical tree diagram illustrating cell type relationships across 45 human tissue datasets. **b,** UMAP visualizations of cell types from 30 human tissues not included in Fig. 6e.

**Extended Data Fig.18 | Hierarchical Structure and Visualization of Cell Types in the Mouse Tissue Atlas. a,** Hierarchical tree diagram illustrating cell type relationships across 23 mouse tissue datasets. **b,** UMAP visualizations of cell types from 13 mouse tissues not included in Fig. 6g.

**Extended Data Fig. 19 | UniCell Enhances Embedding Quality of Single-Cell Foundation Models. a, c, e,** Bar plots showing Average Silhouette Width (ASW) scores based on cell type labels for scGPT (**a**), GeneFormer (**c**), and scFoundation (**e**), compared to their UniCell-integrated counterparts. Higher ASW scores indicate better separation of cell types in the embedding space. **b, d, f,** UMAP visualizations of 45 human datasets based on embeddings from scGPT (**b**), GeneFormer (**d**), and scFoundation (**f**), with and without UniCell integration. Colors represent different cell types.

**Extended Data Fig.20 | UniCell Enhances Annotation Performance of scGPT Across Hierarchical Cell Types. a,** Scatter plot comparing ASW scores between scGPT and scGPT+UniCell across 45 human tissue datasets. Each dot represents one dataset; improved ASW reflects better-defined cell type clusters. **b,** Bubble plot showing differences in annotation Accuracy and Macro-F1 between scGPT and scGPT+UniCell. Dot size corresponds to the number of cells in each tissue. **c,** Line charts comparing Macro-F1 scores between scGPT and scGPT+UniCell across cell types at different hierarchical levels, demonstrating consistent performance gains from UniCell integration.

## Notes

### Competing Interest Statement

The authors have declared no competing interest.

