## Supplementary figures and images for "UniCell: Towards a Unified Solution for Cell Annotation, Nomenclature Harmonization, Atlas Construction in Single-Cell Transcriptomics"

### Supplemental Figure 1

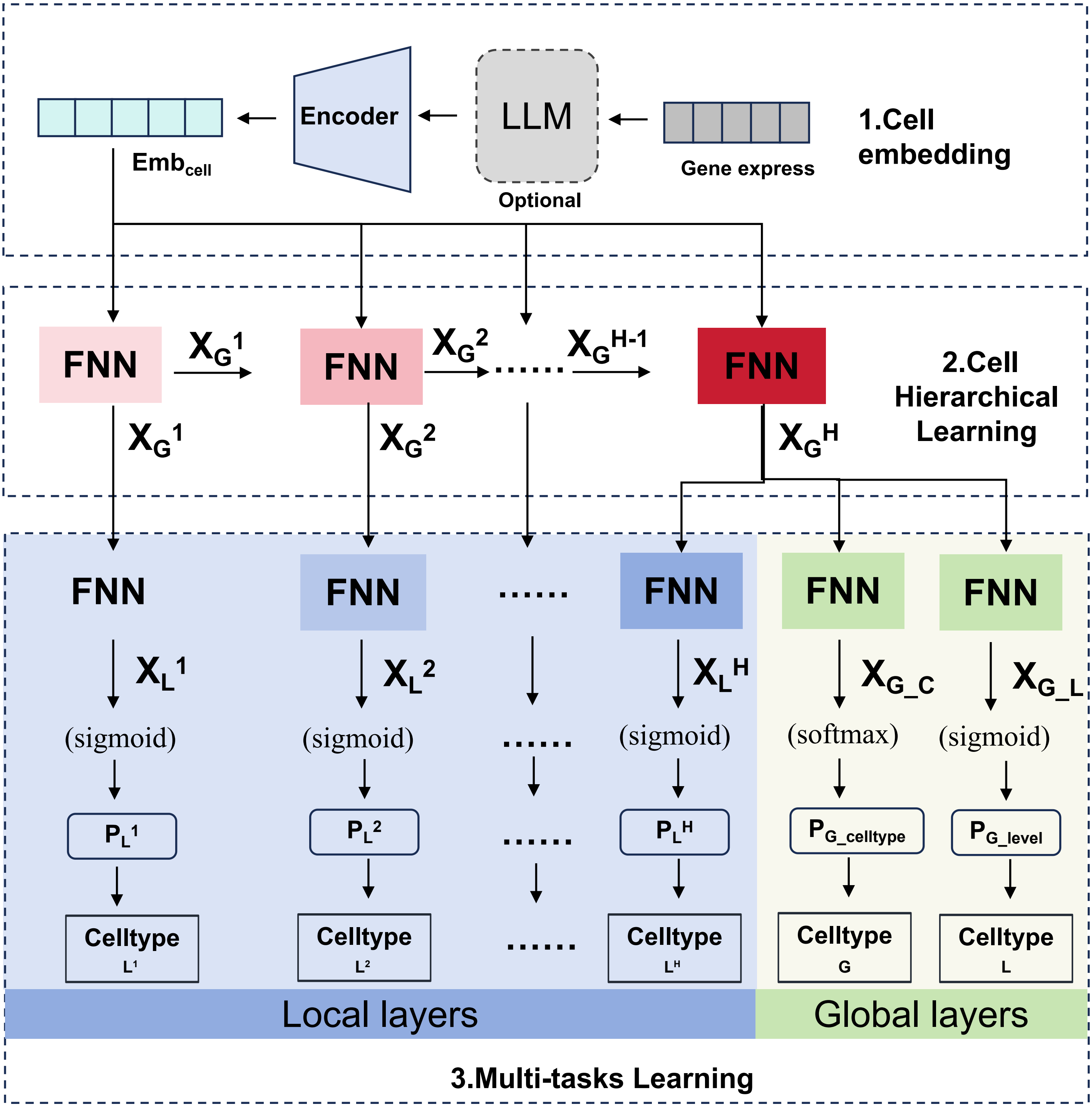

### Supplemental Figure 2

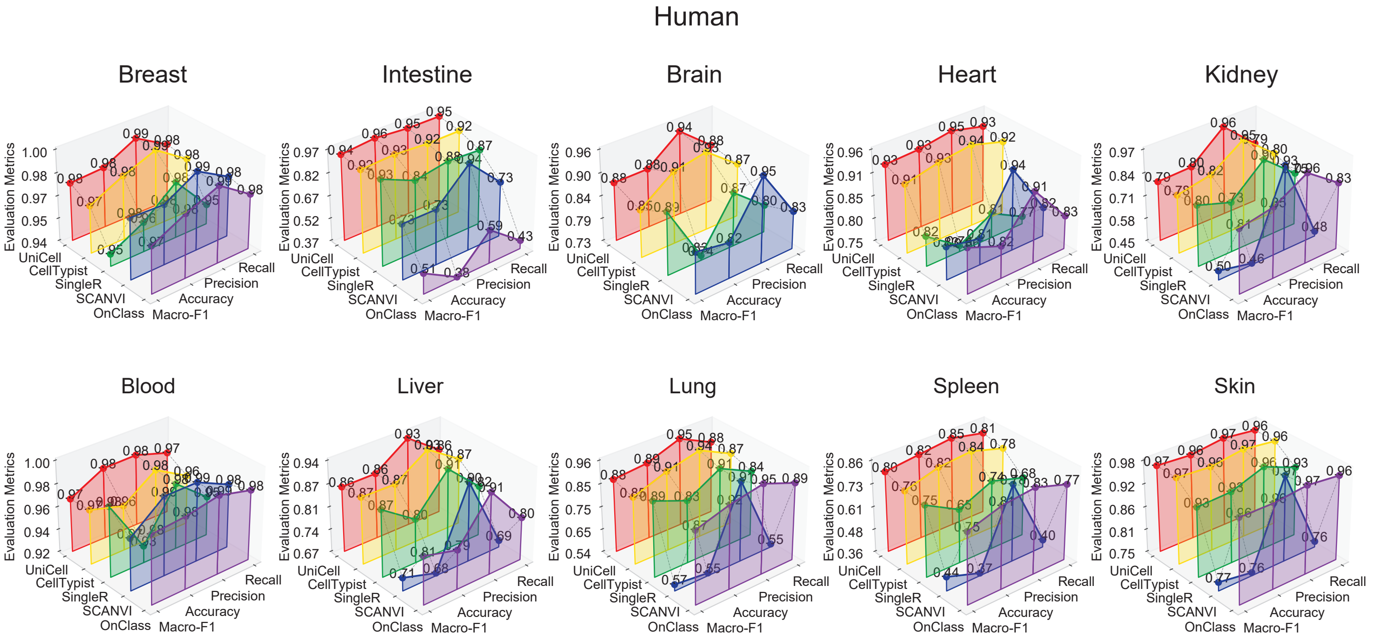

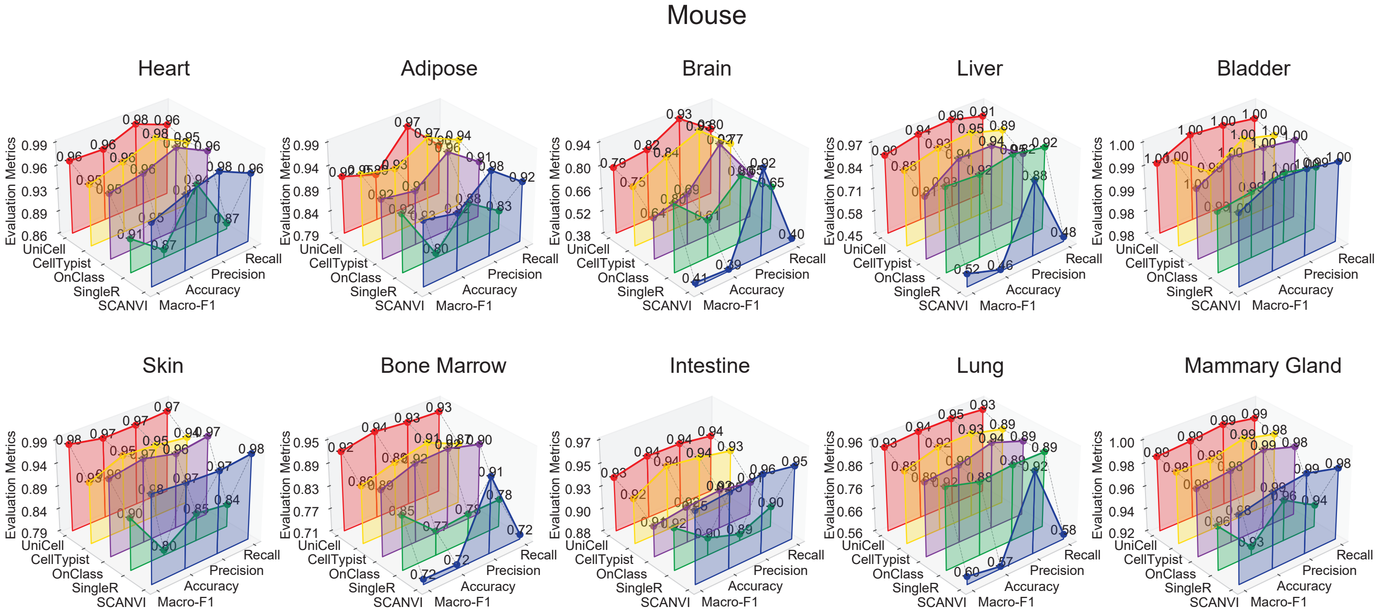

### Supplemental Figure 3

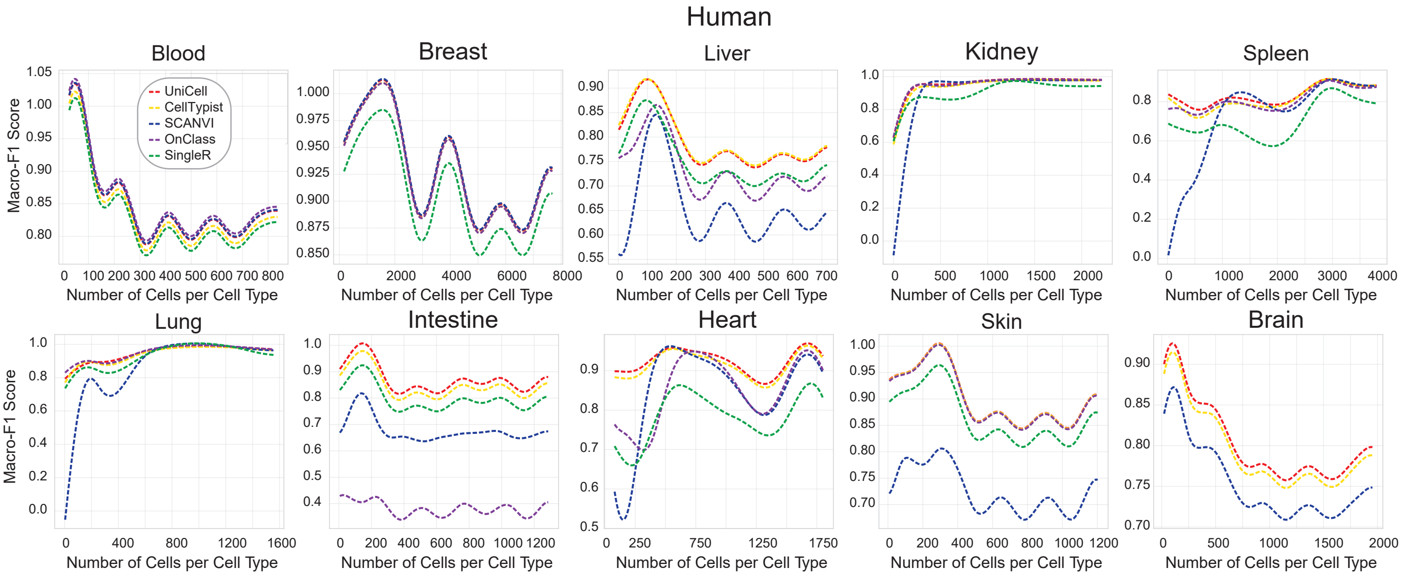

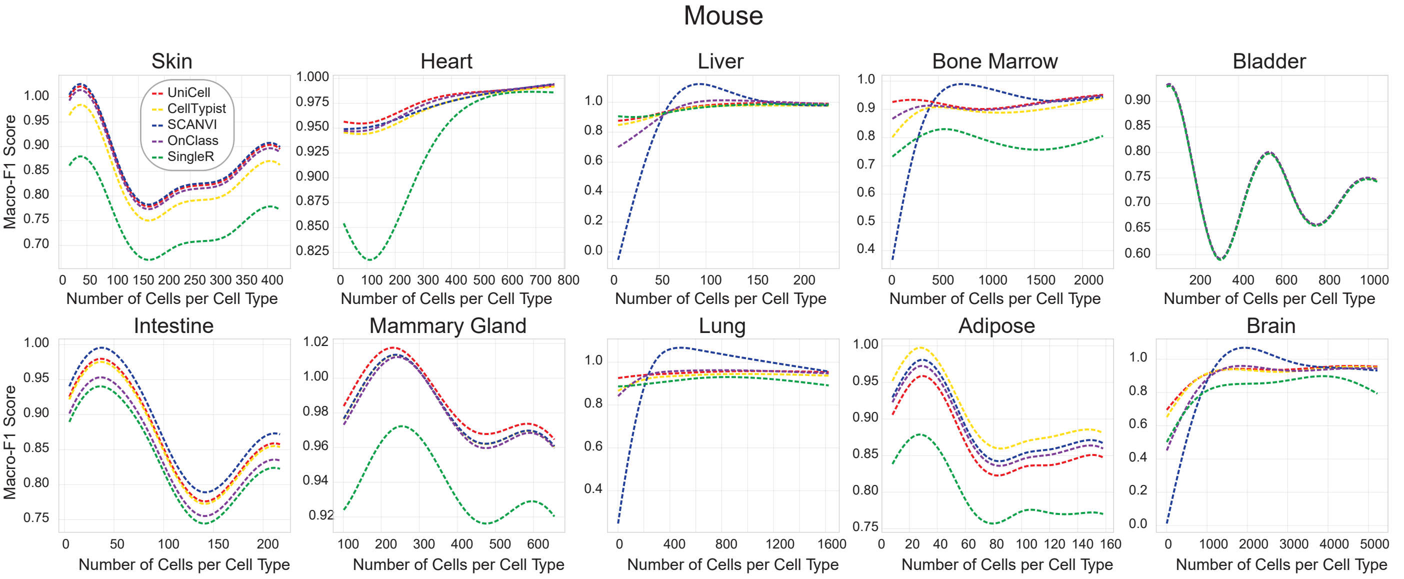

### Supplemental Figure 4

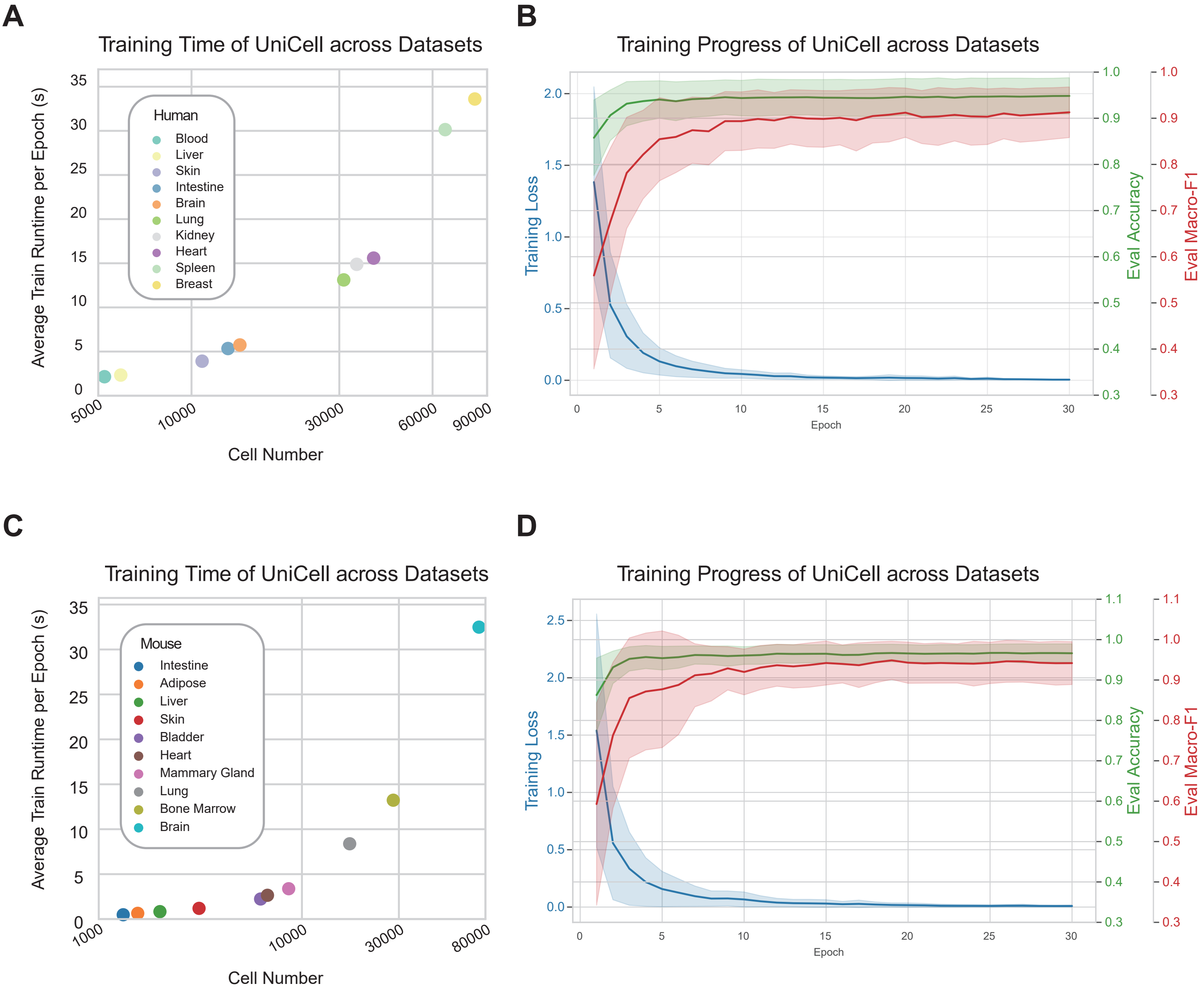

### Supplemental Figure 5

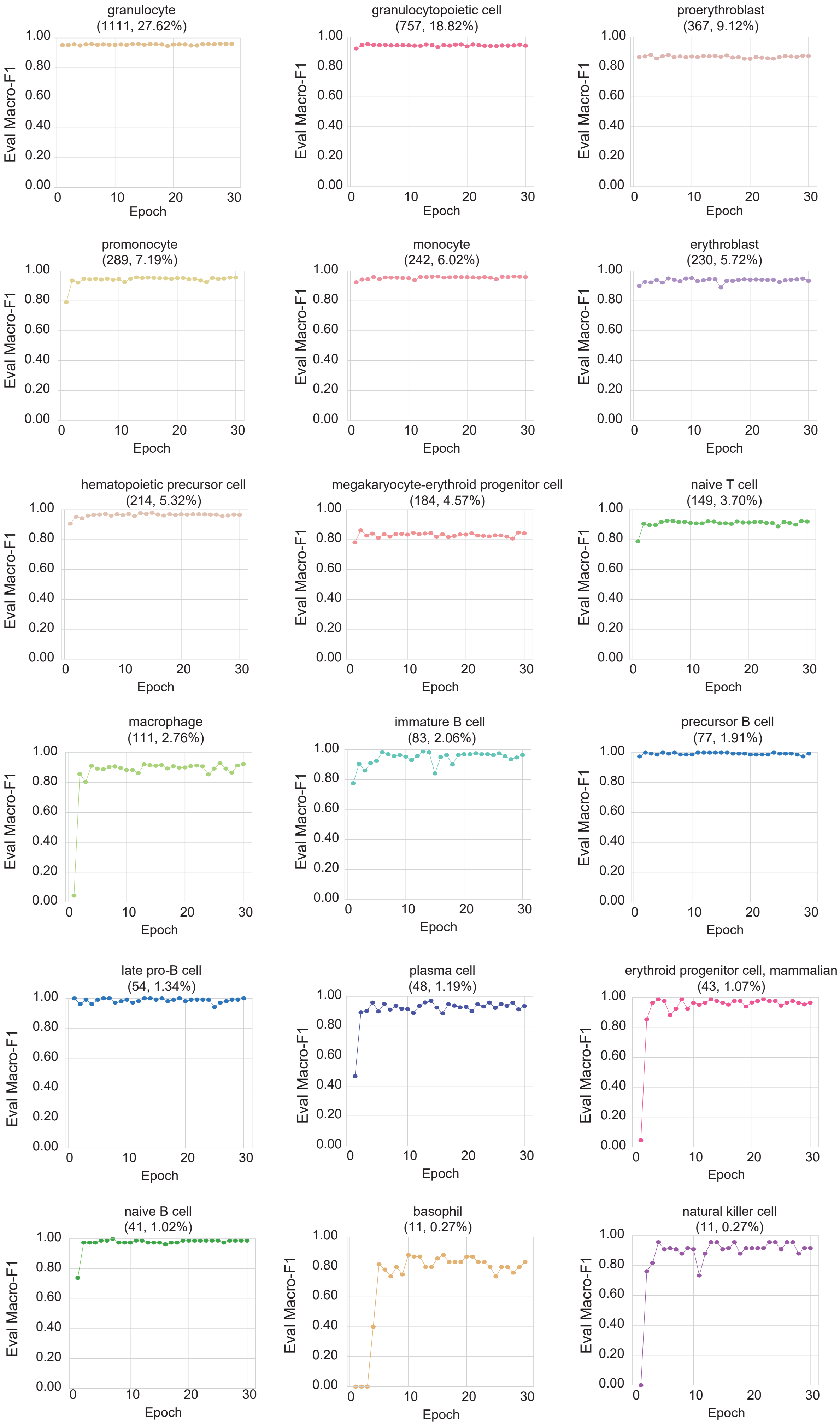

### Supplemental Figure 6

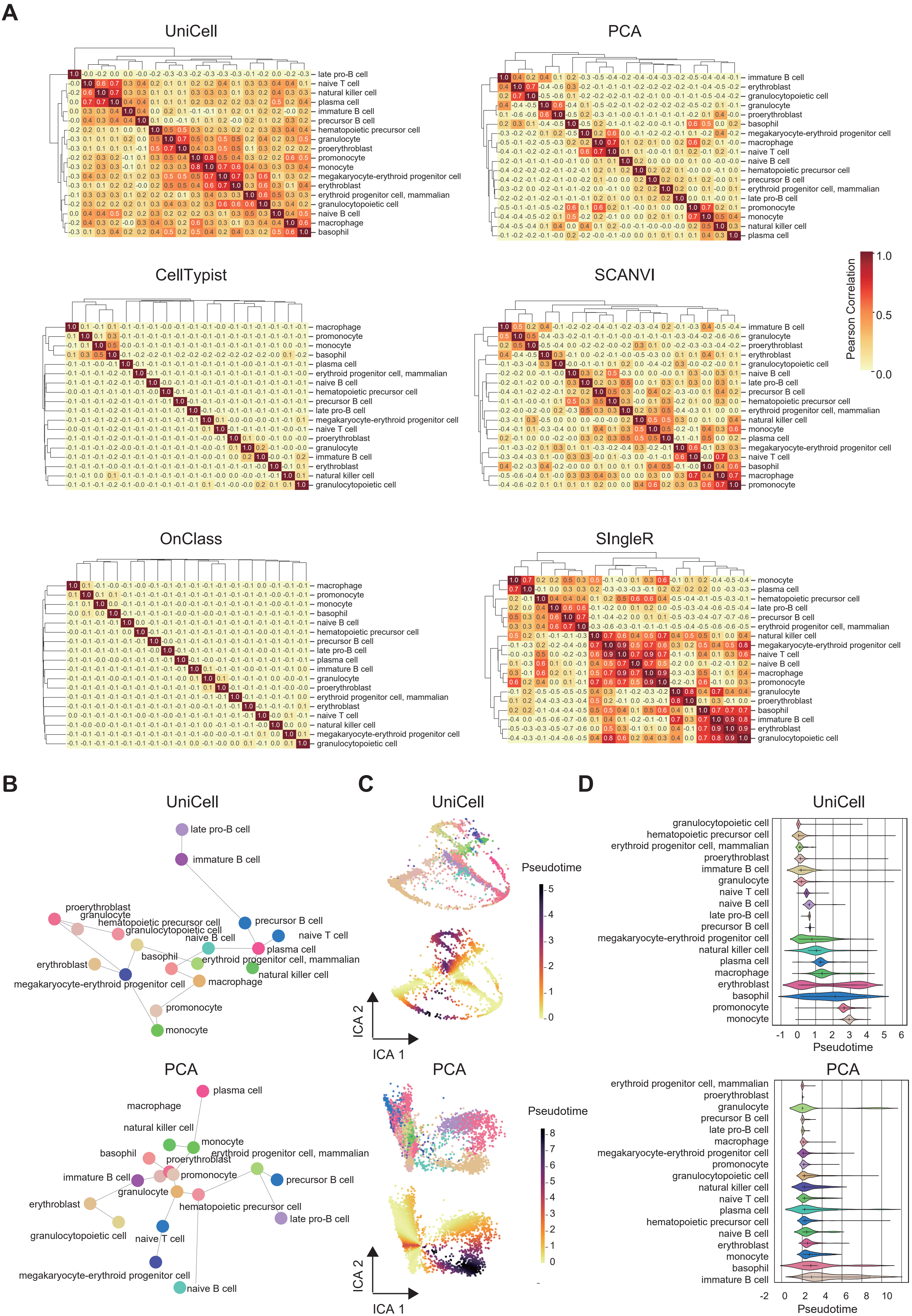

### Supplemental Figure 7

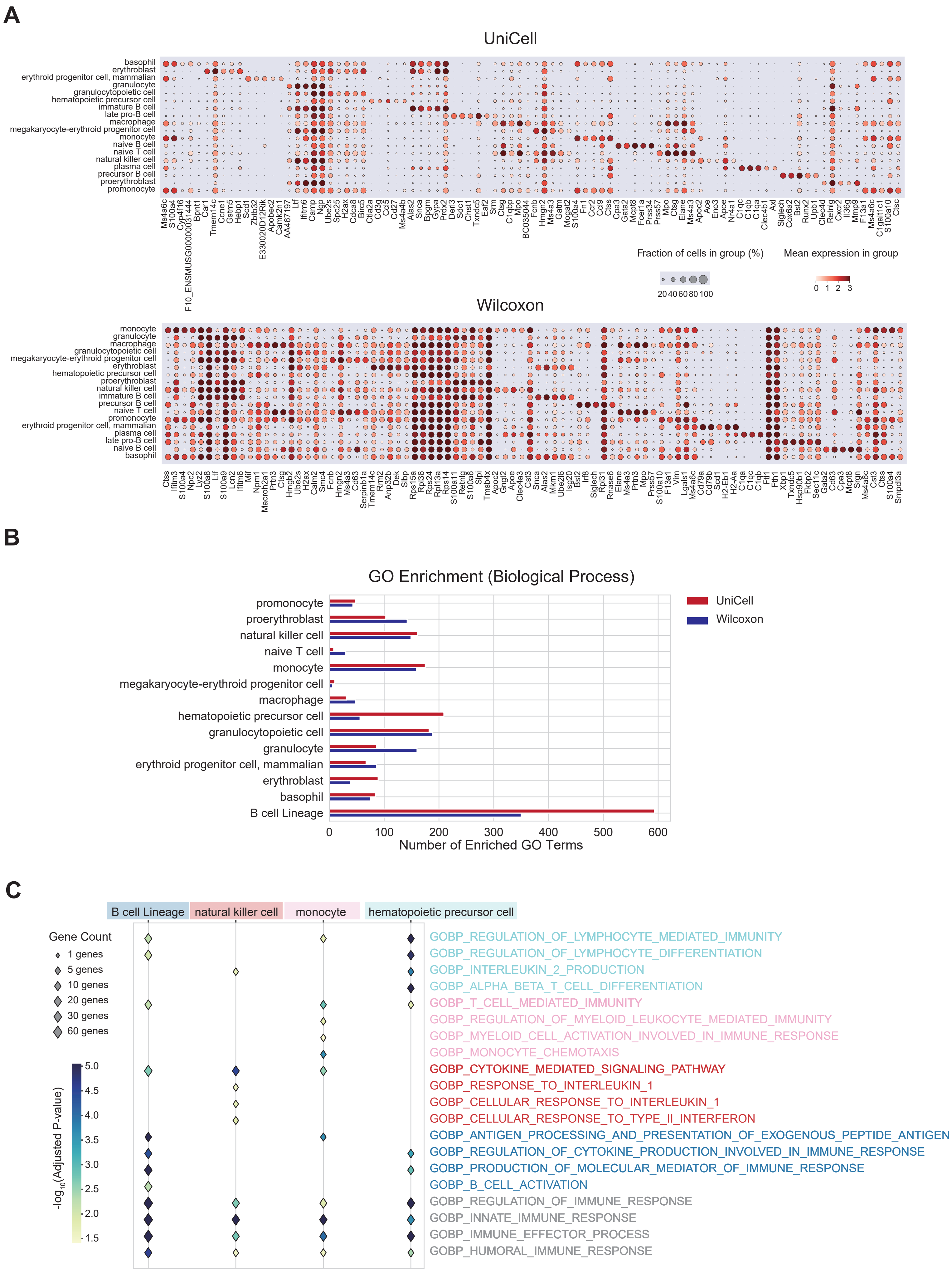

### Supplemental Figure 8

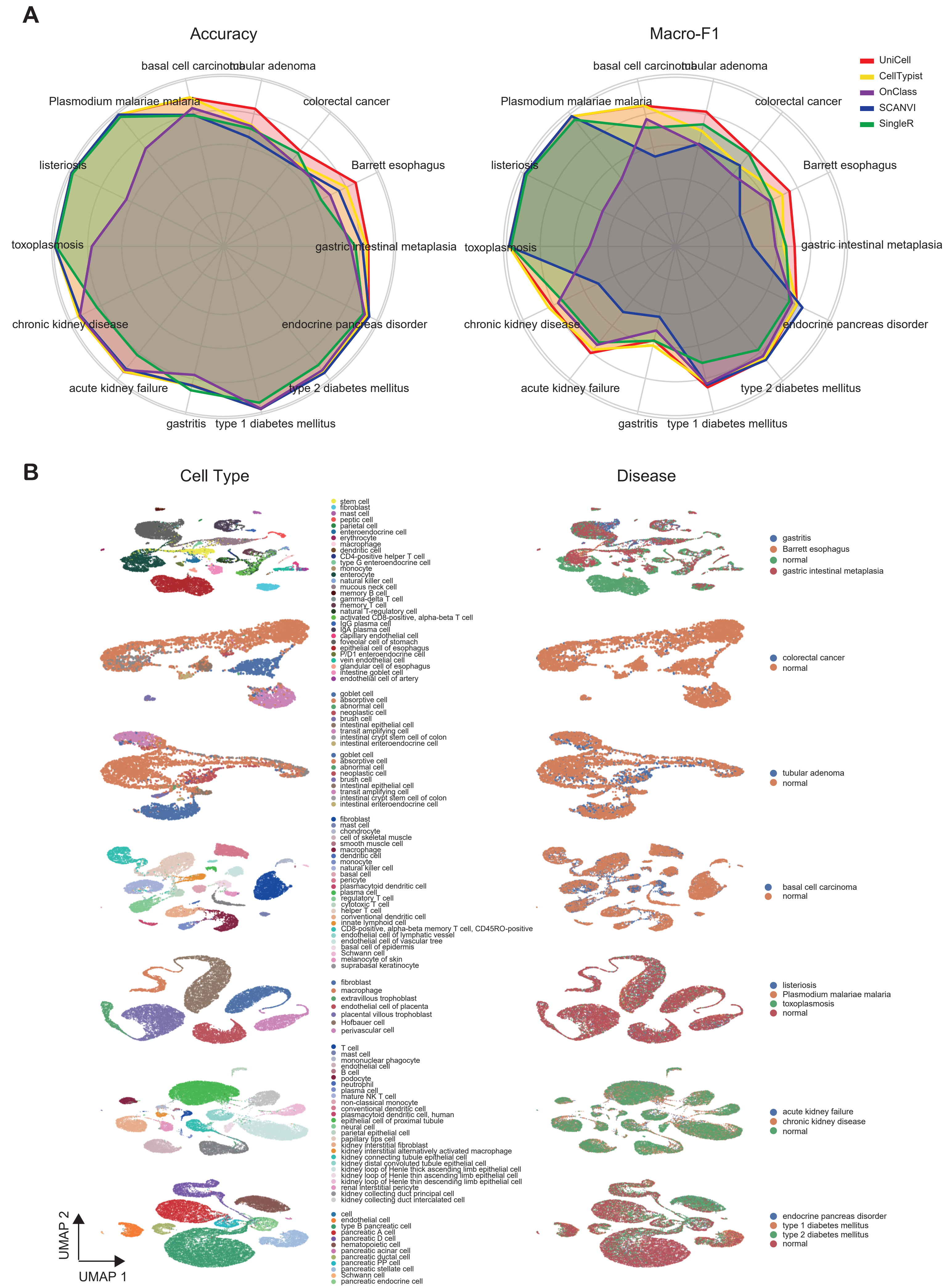

### Supplemental Figure 9

#### Figure S9

## A

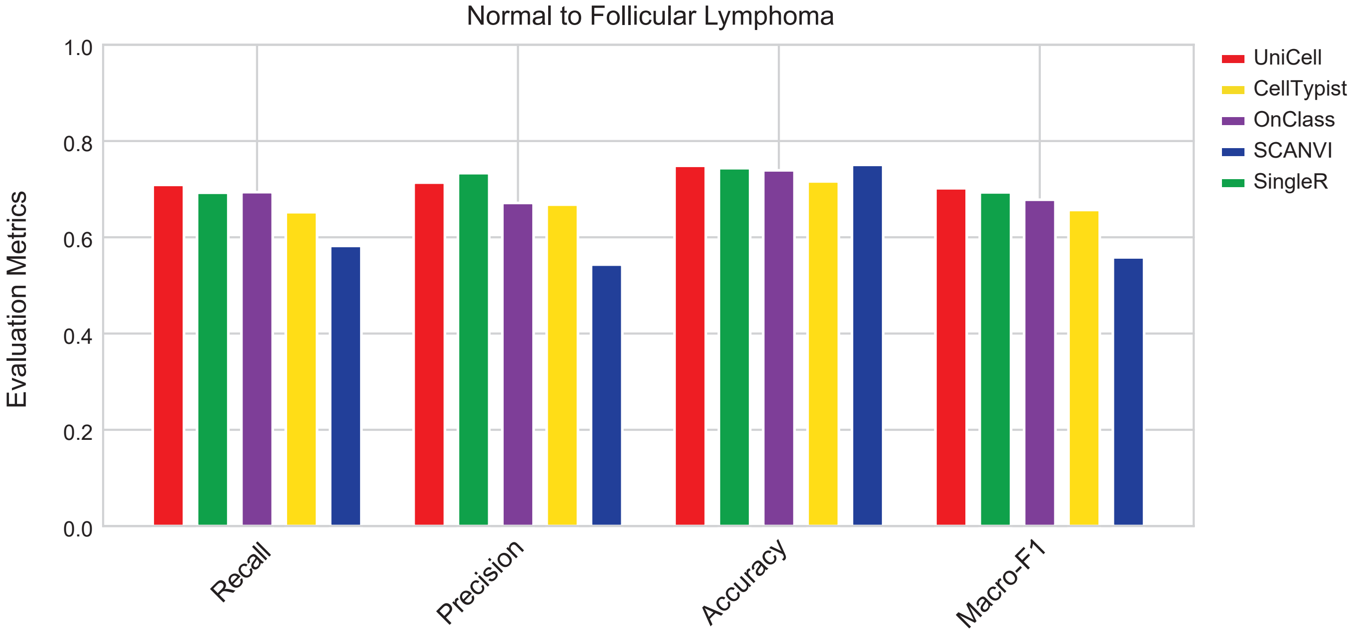


## B

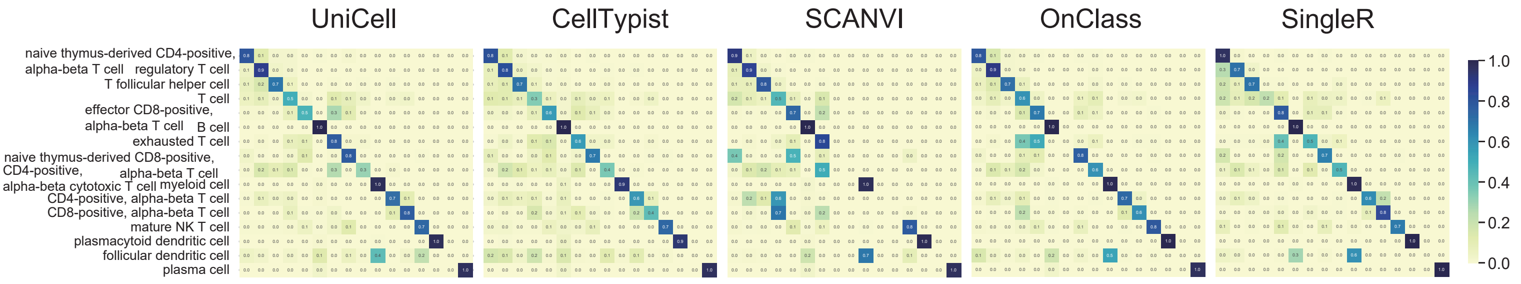


## C

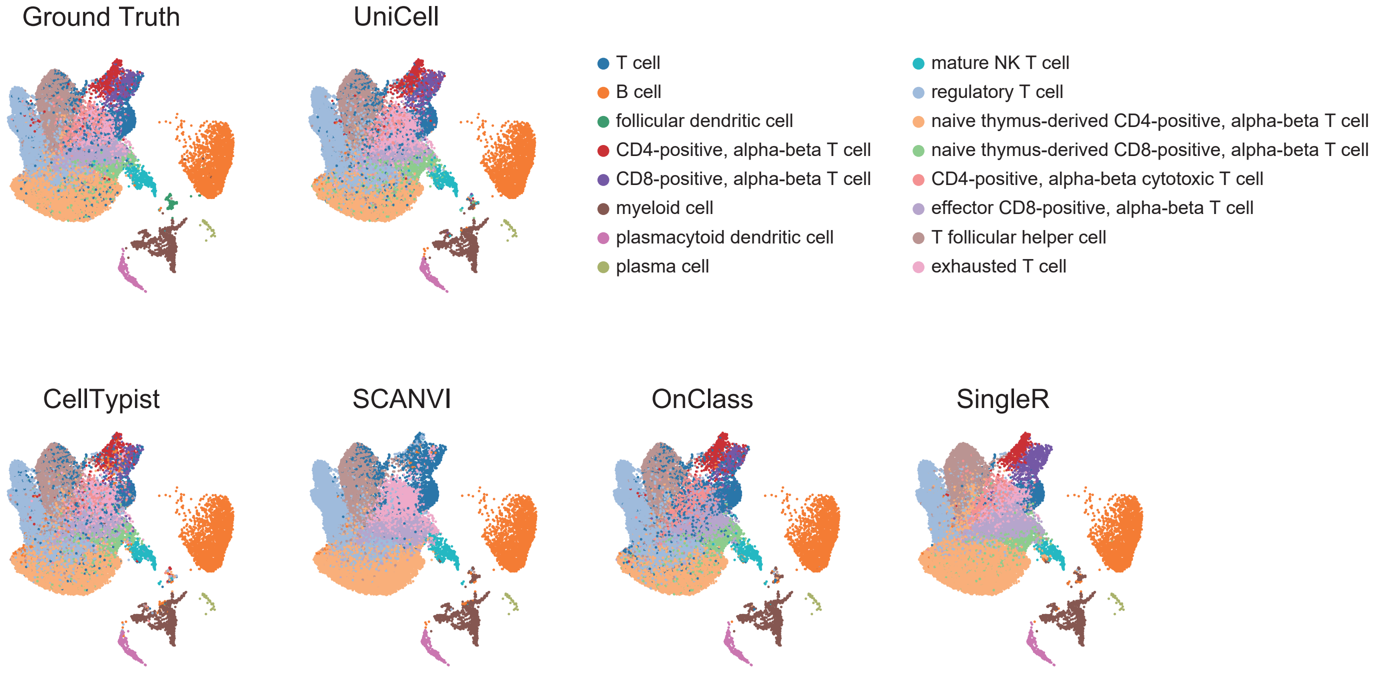

### Supplemental Figure 10

#### Figure S10

## A

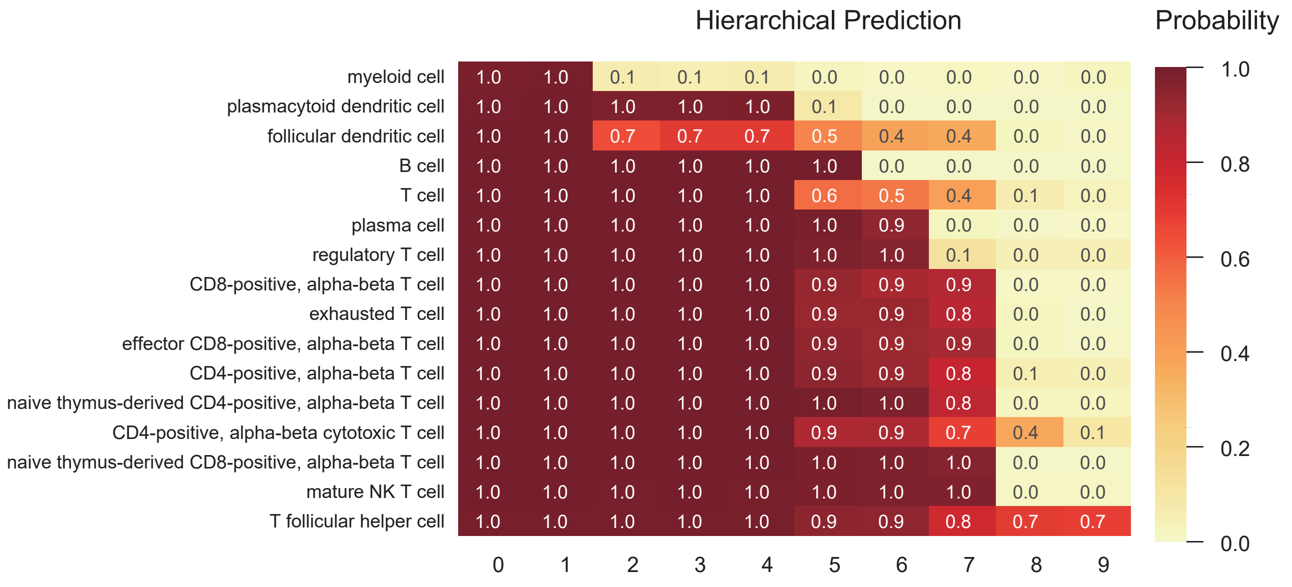


## B

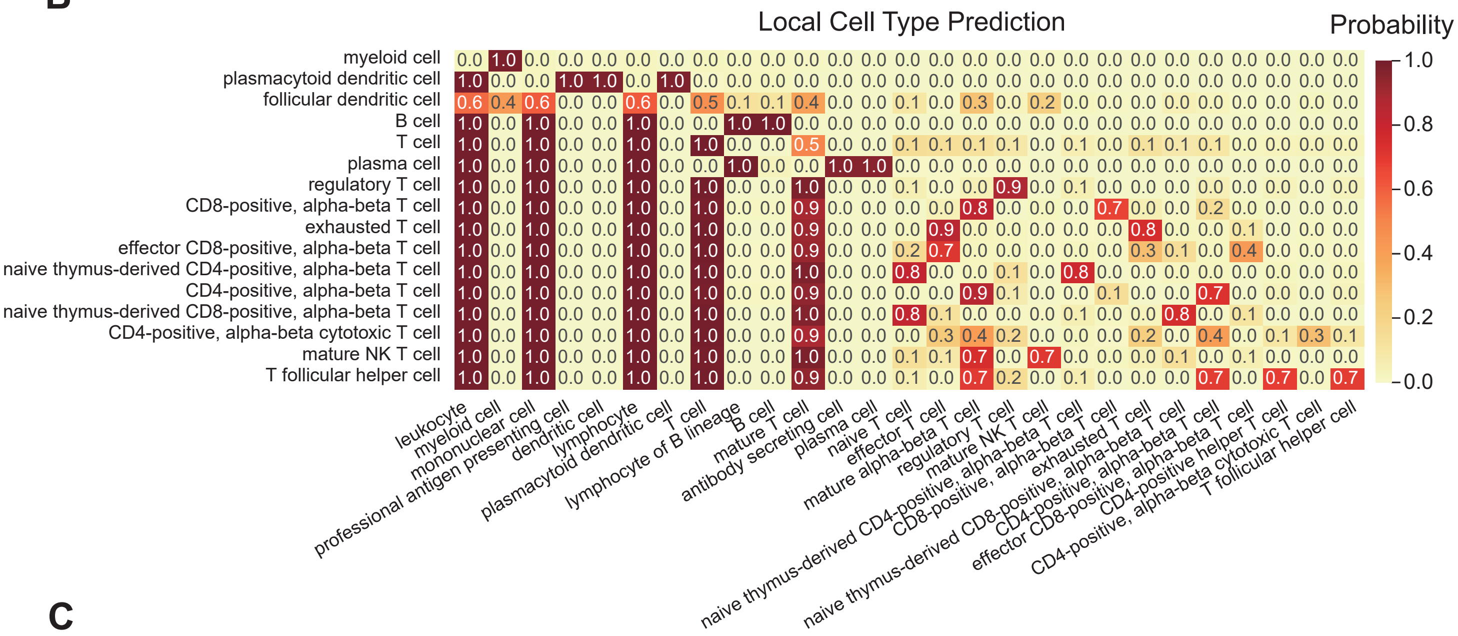


## C

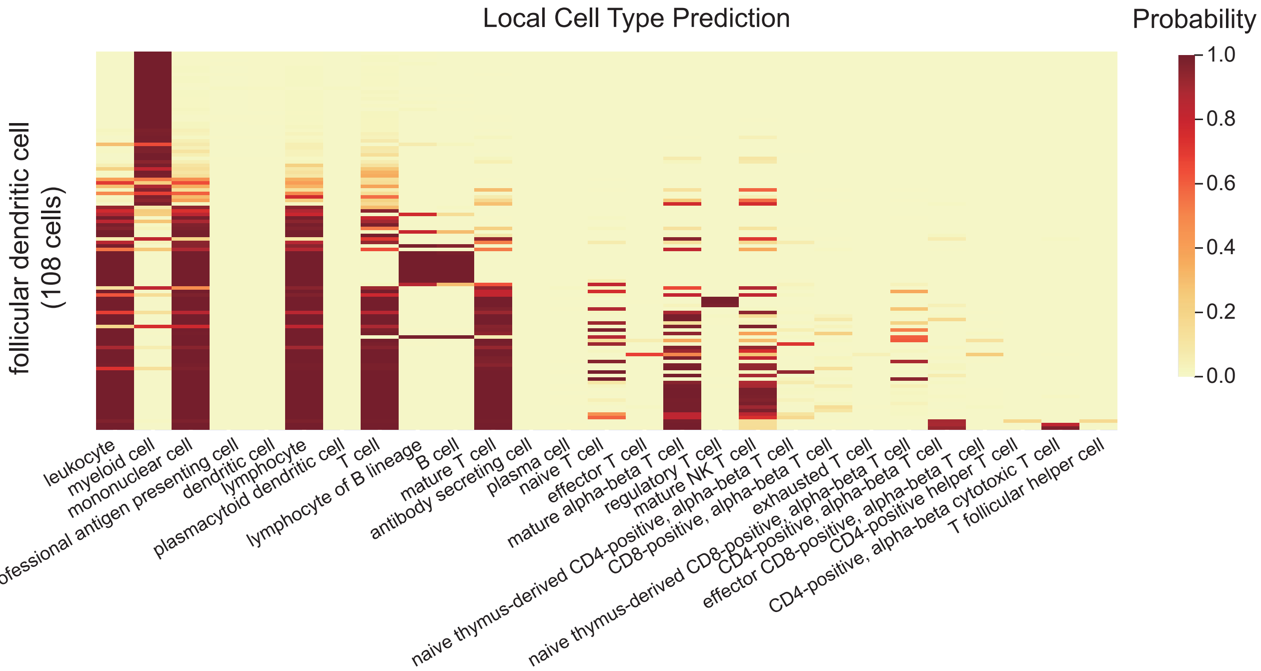

### Supplemental Figure 11

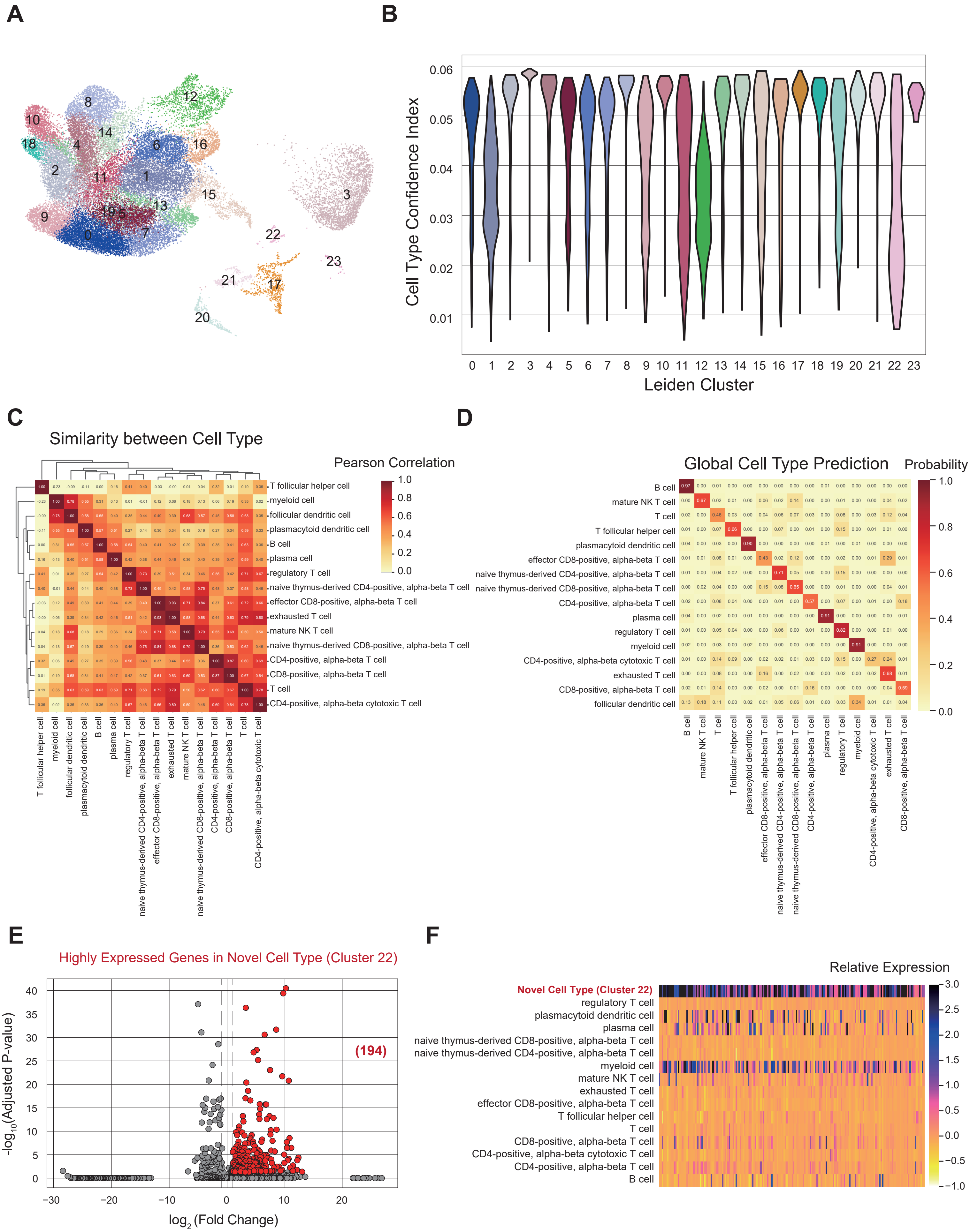

### Supplemental Figure 12

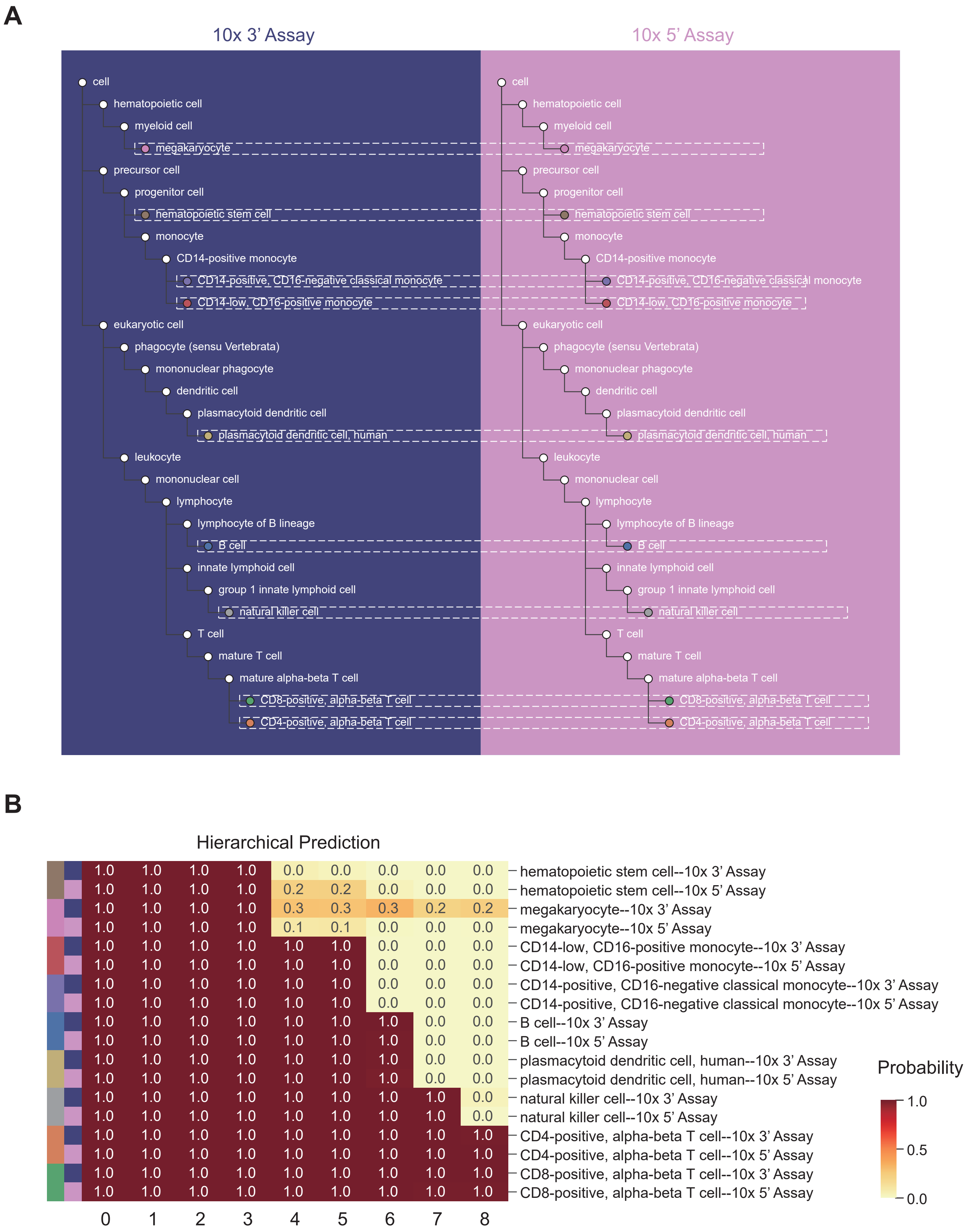

### Supplemental Figure 13

#### Figure S13

**A**

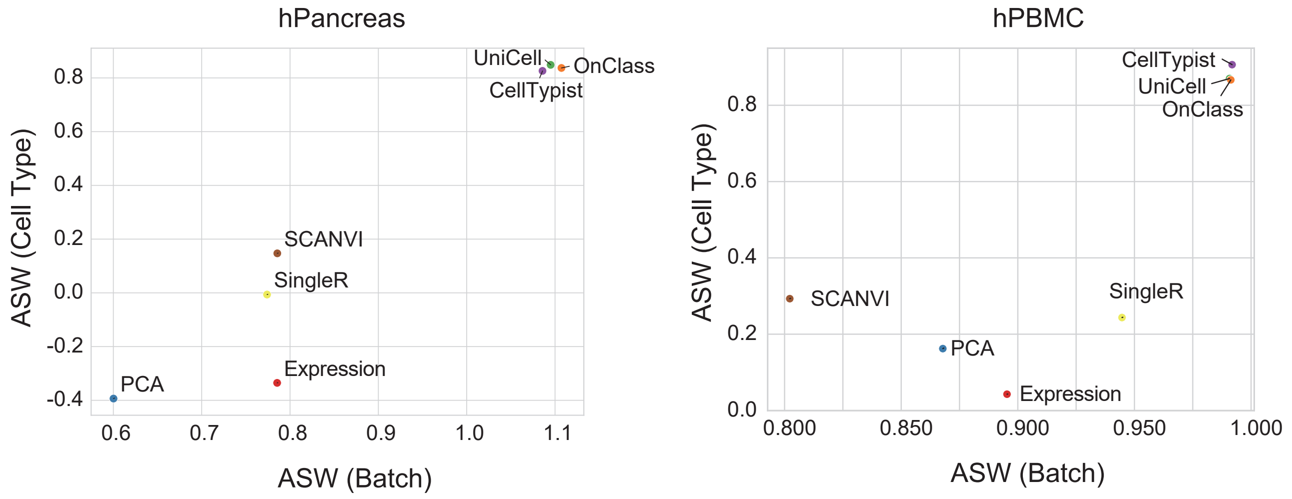


**B**

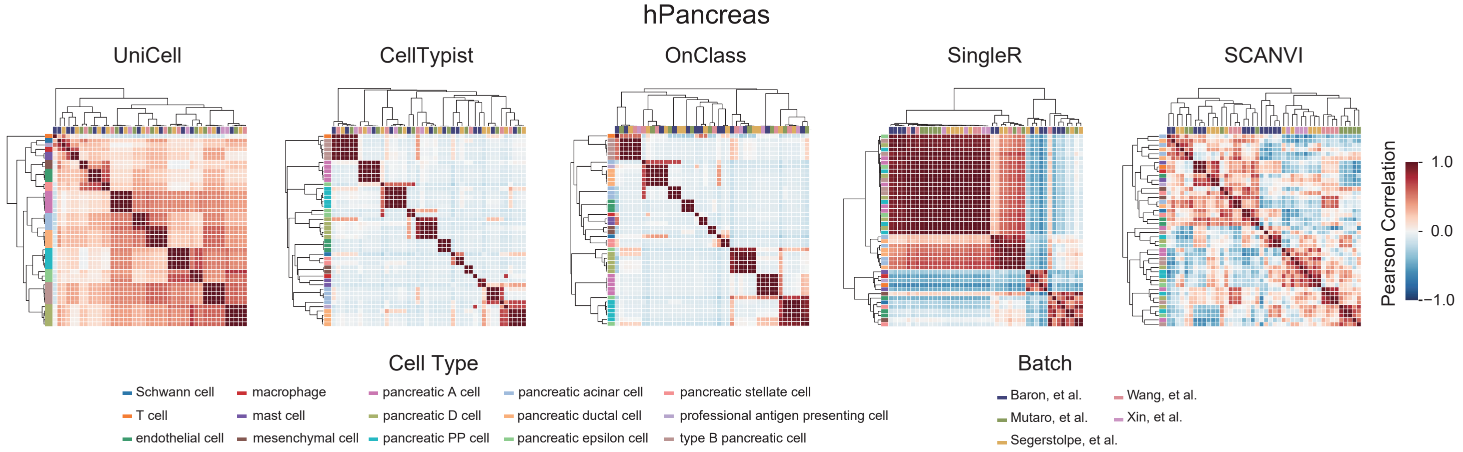


**C**

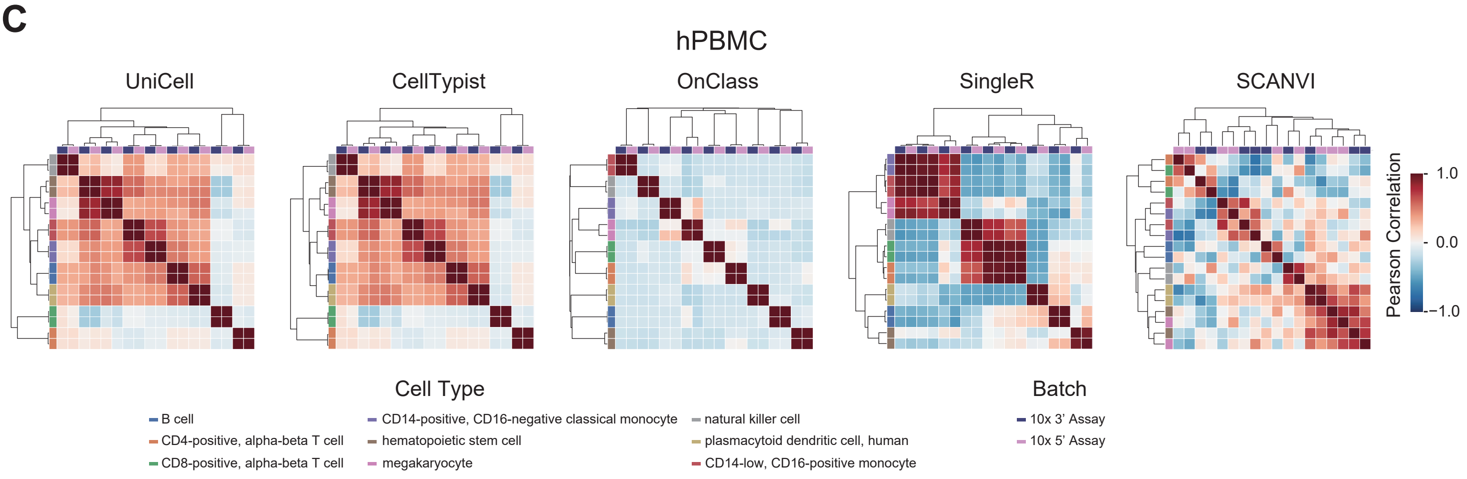


**D**

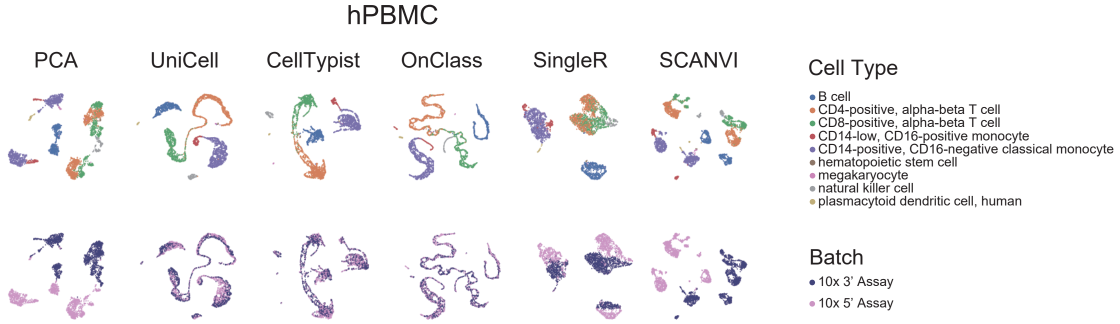

### Supplemental Figure 14

##### Figure S14

A

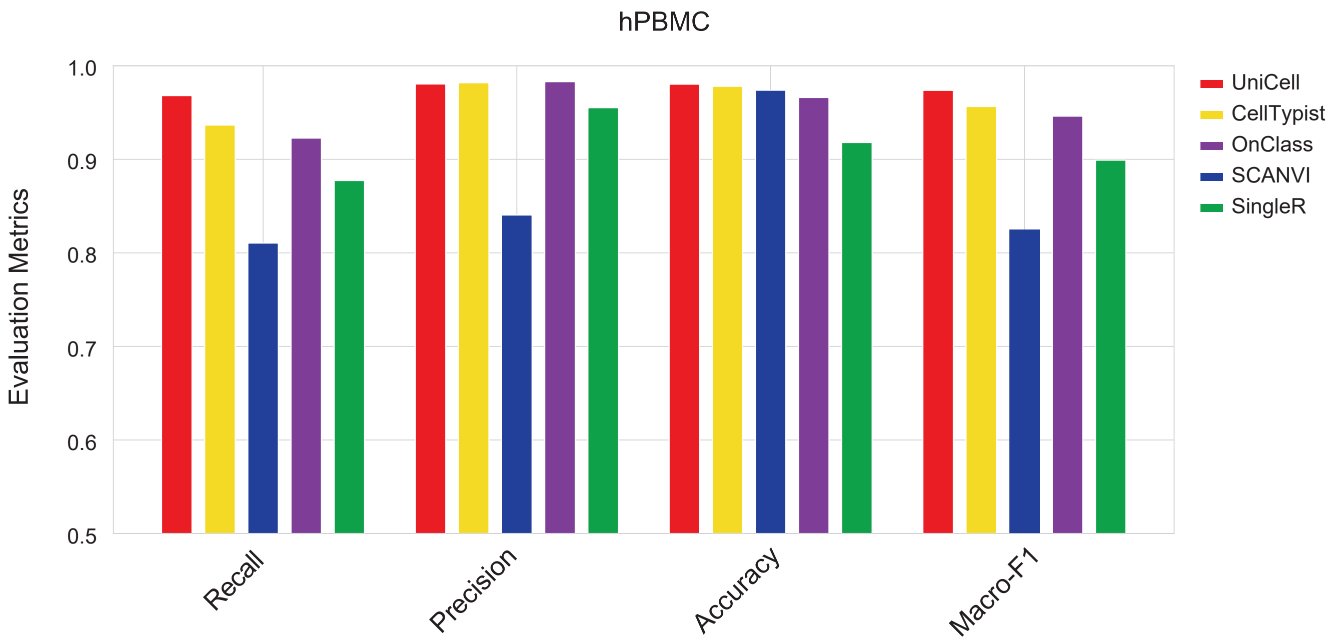


B

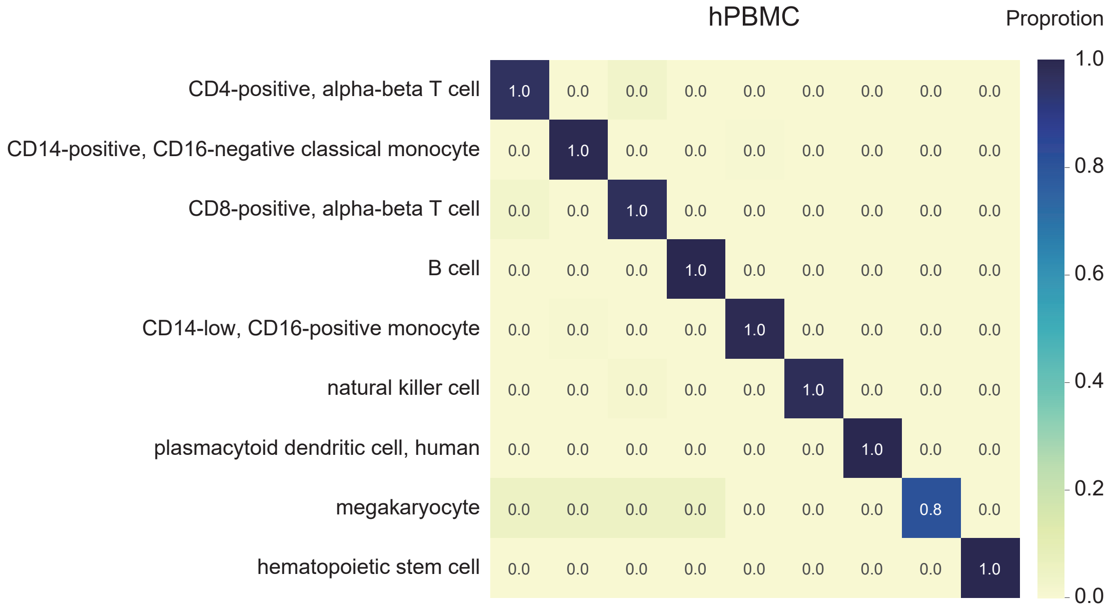

### Supplemental Figure 15

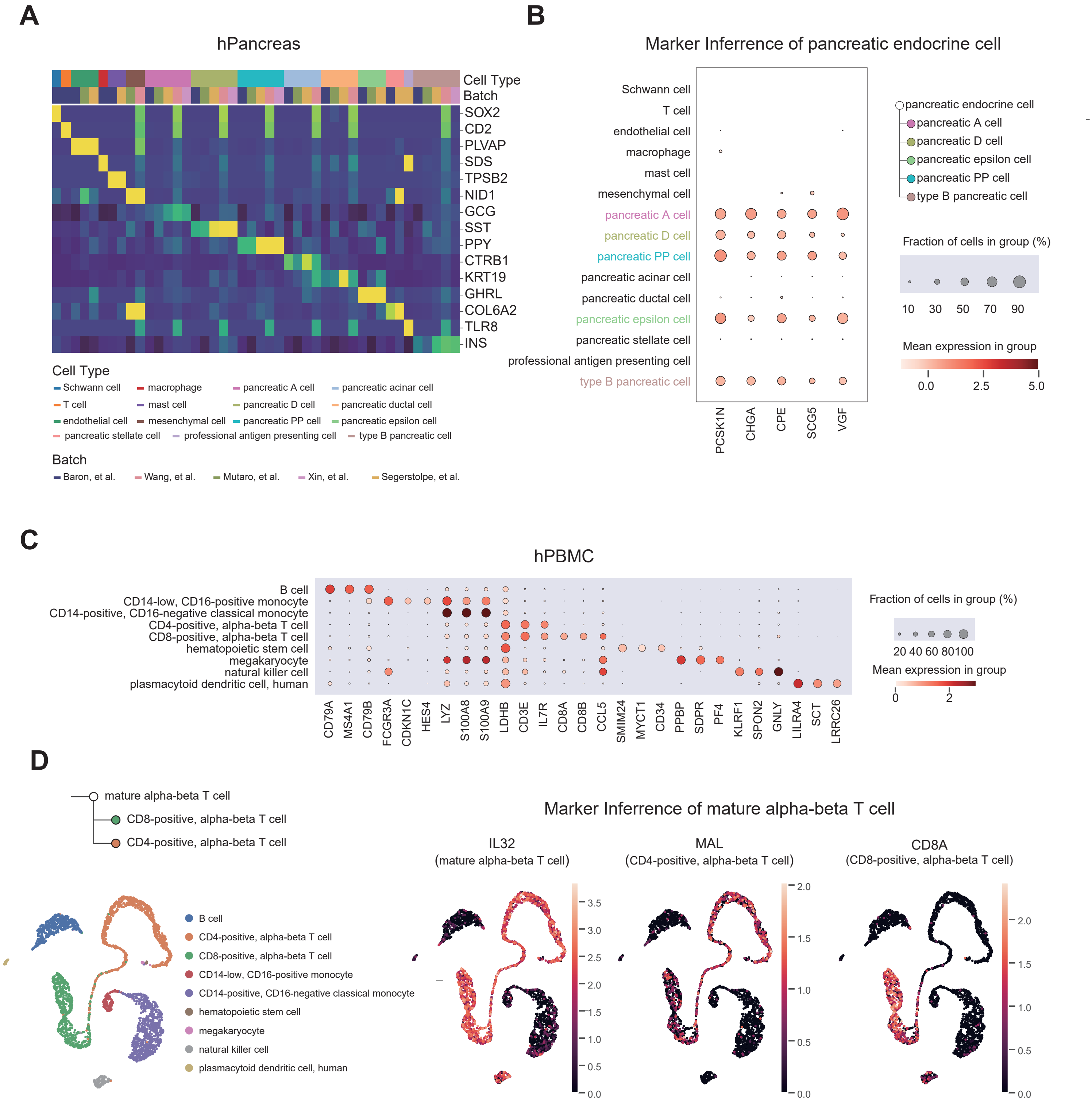

### Supplemental Figure 16

#### Figure S16

## A

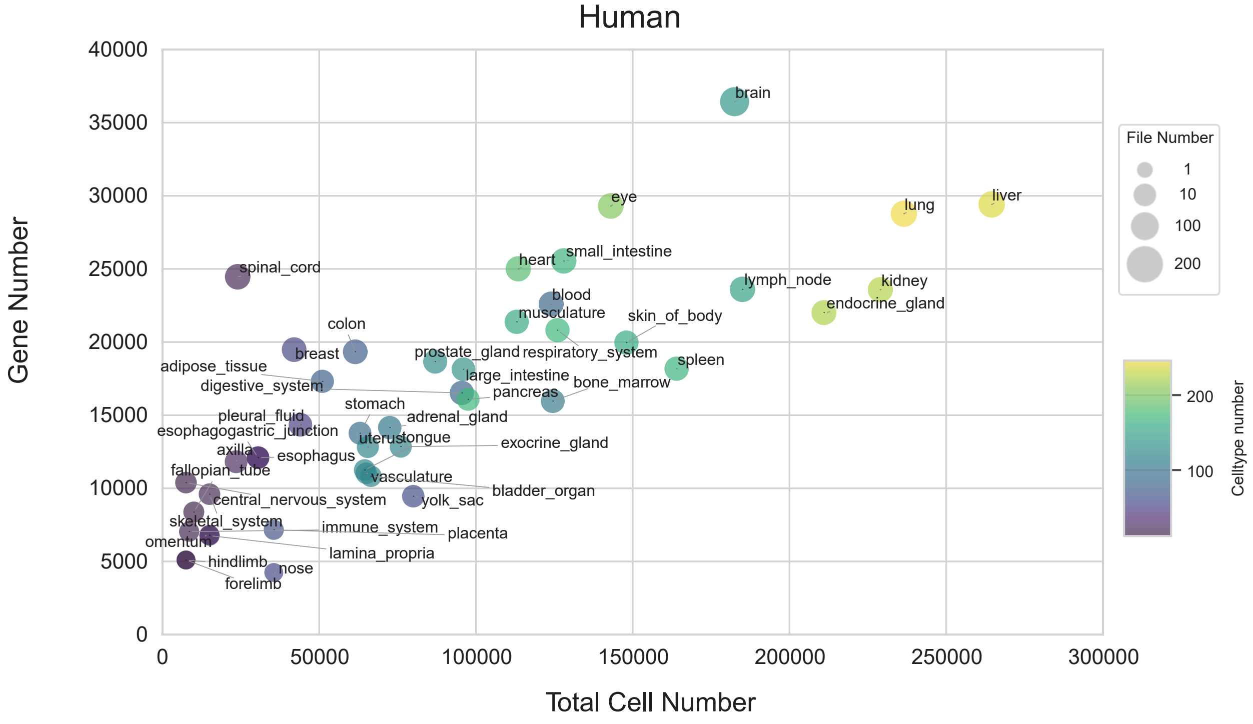


## B

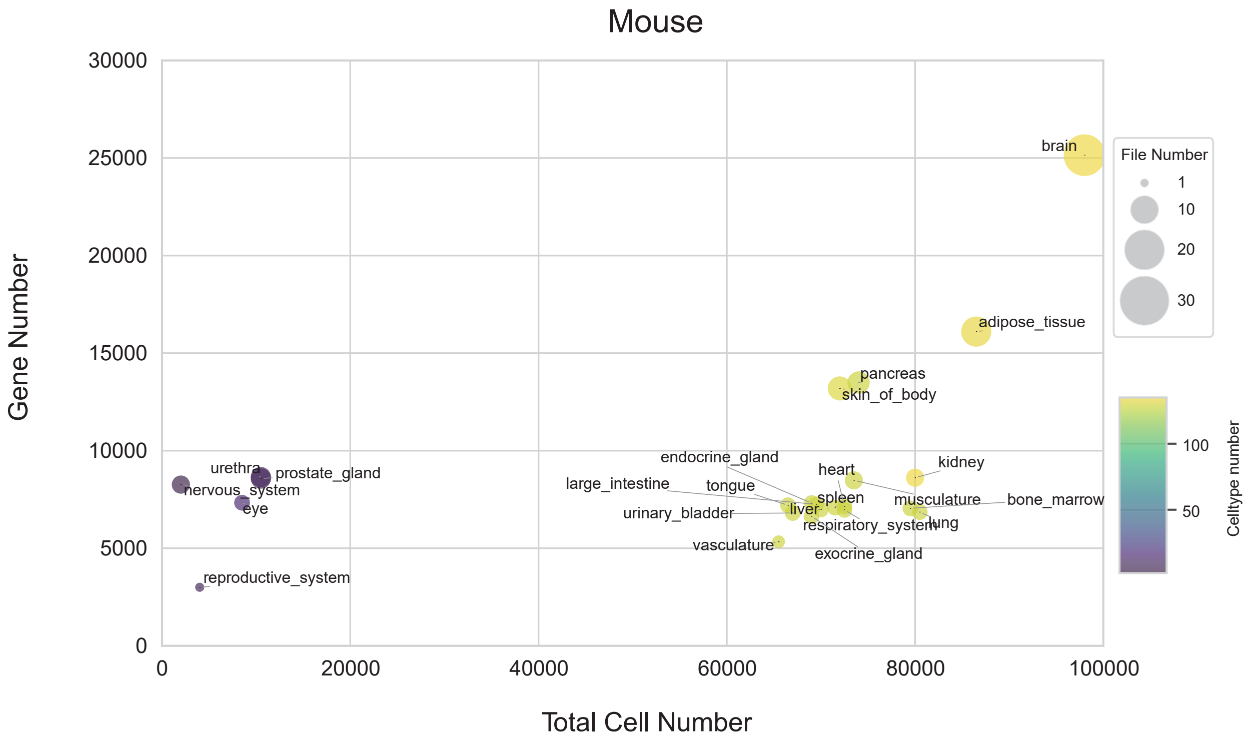

### Supplemental Figure 17

#### Figure S17

## A

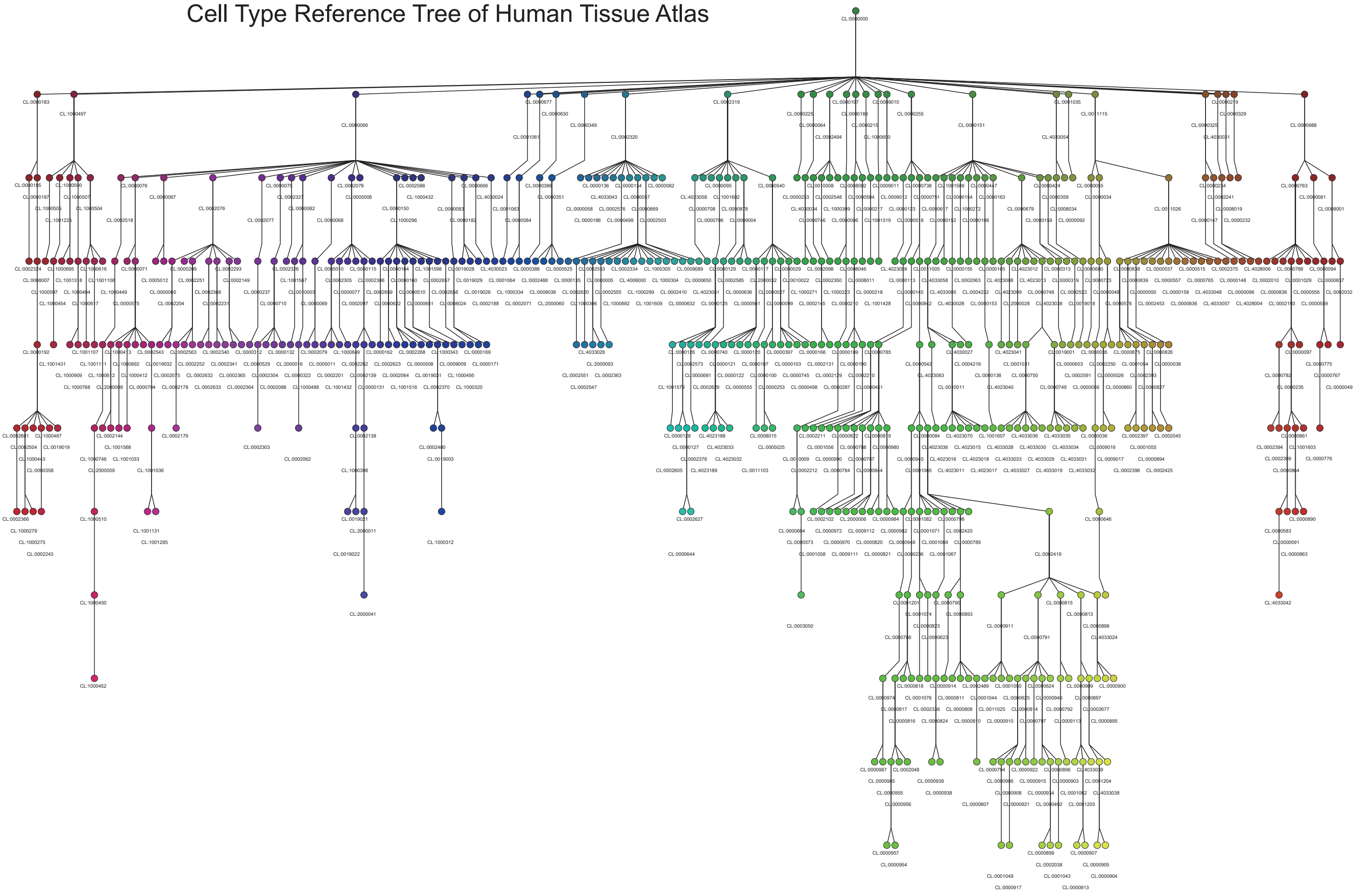


## B

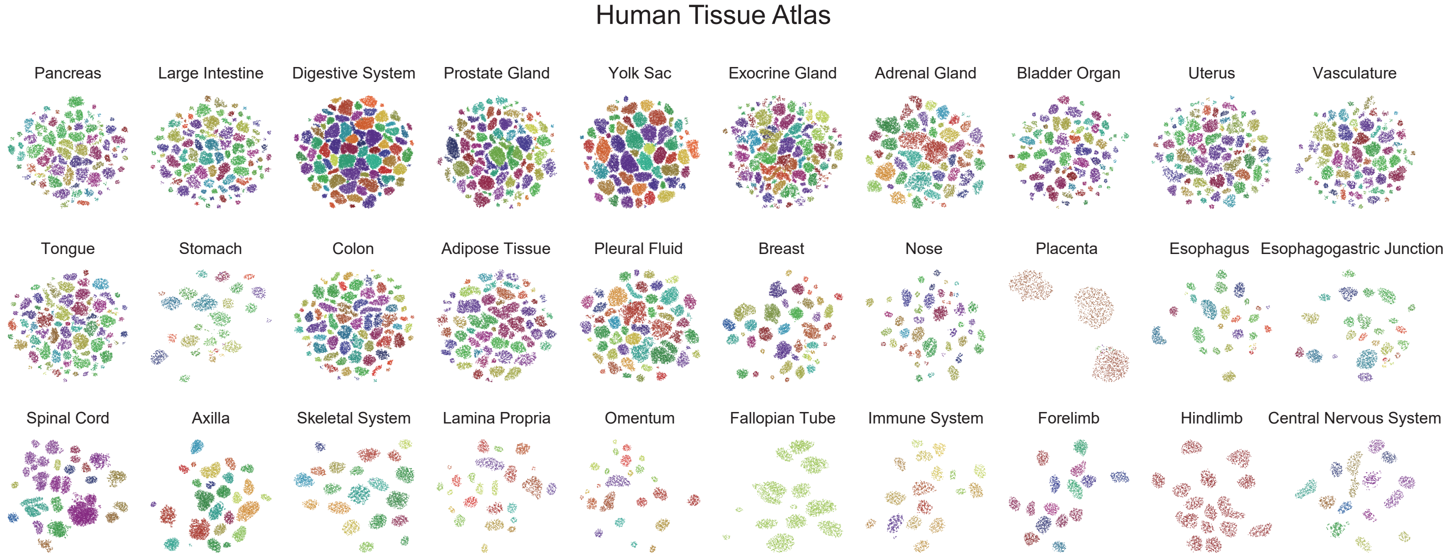

### Supplemental Figure 18

#### Figure S18

**A**

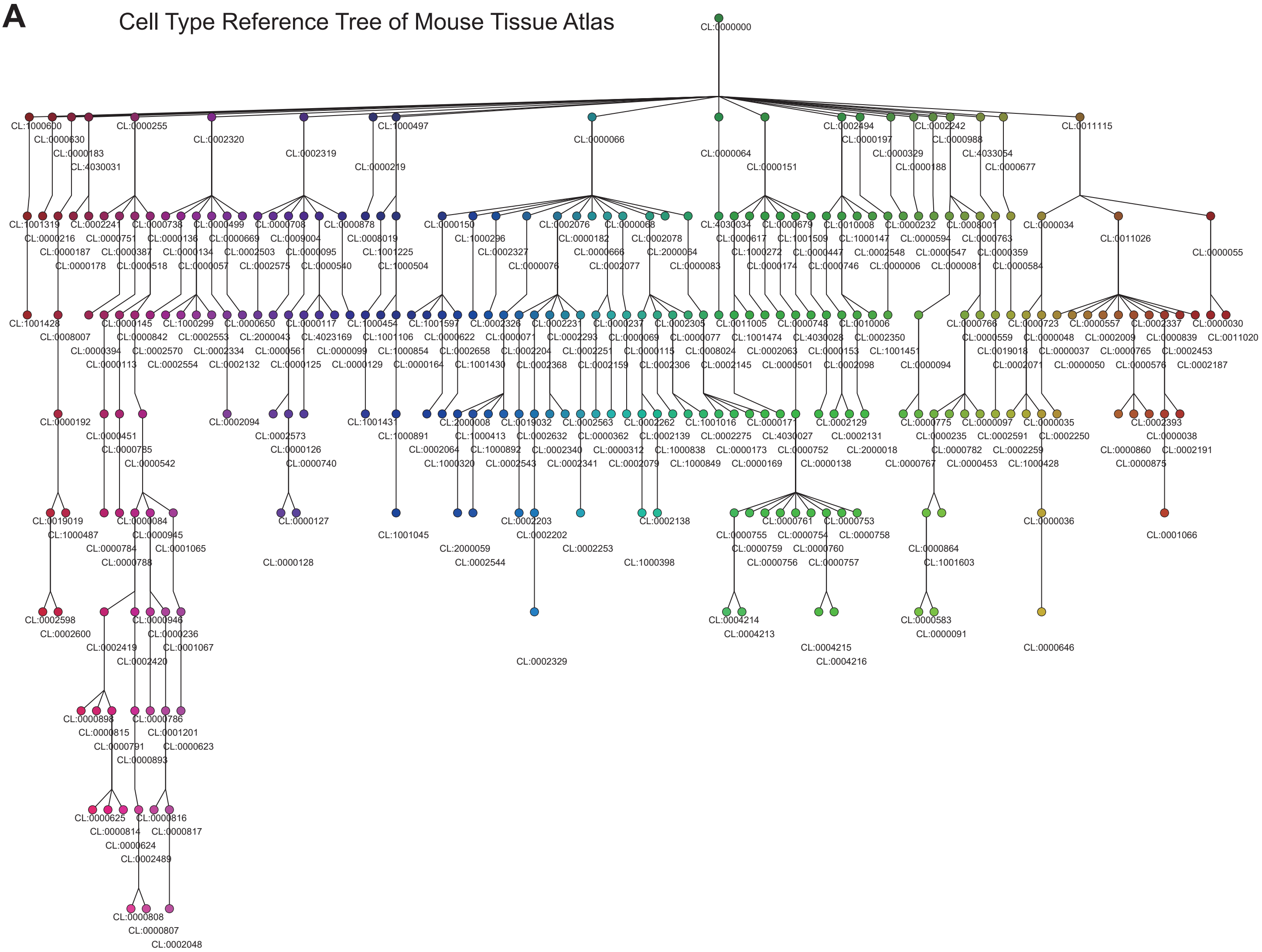


**B**

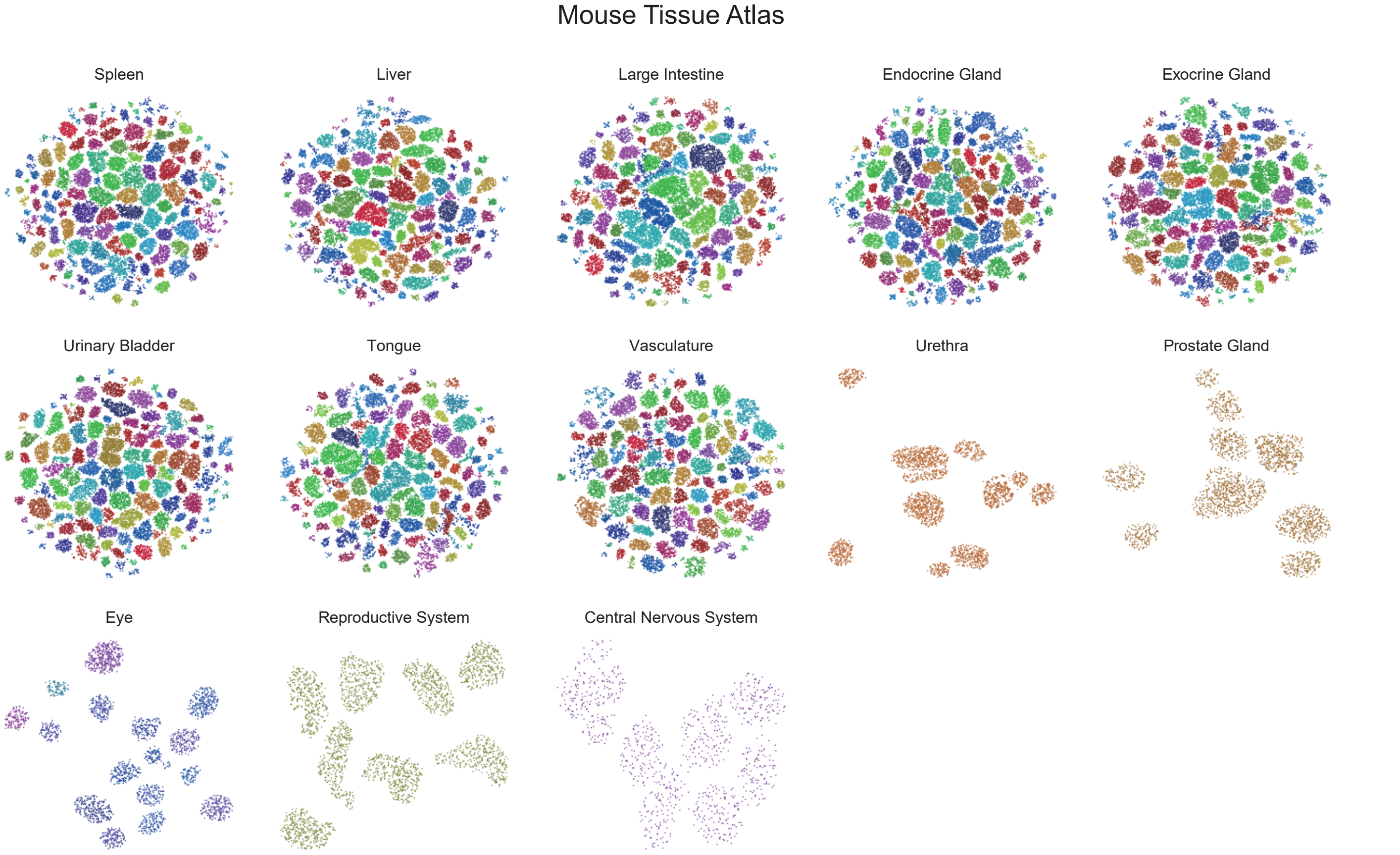

### Supplemental Figure 19

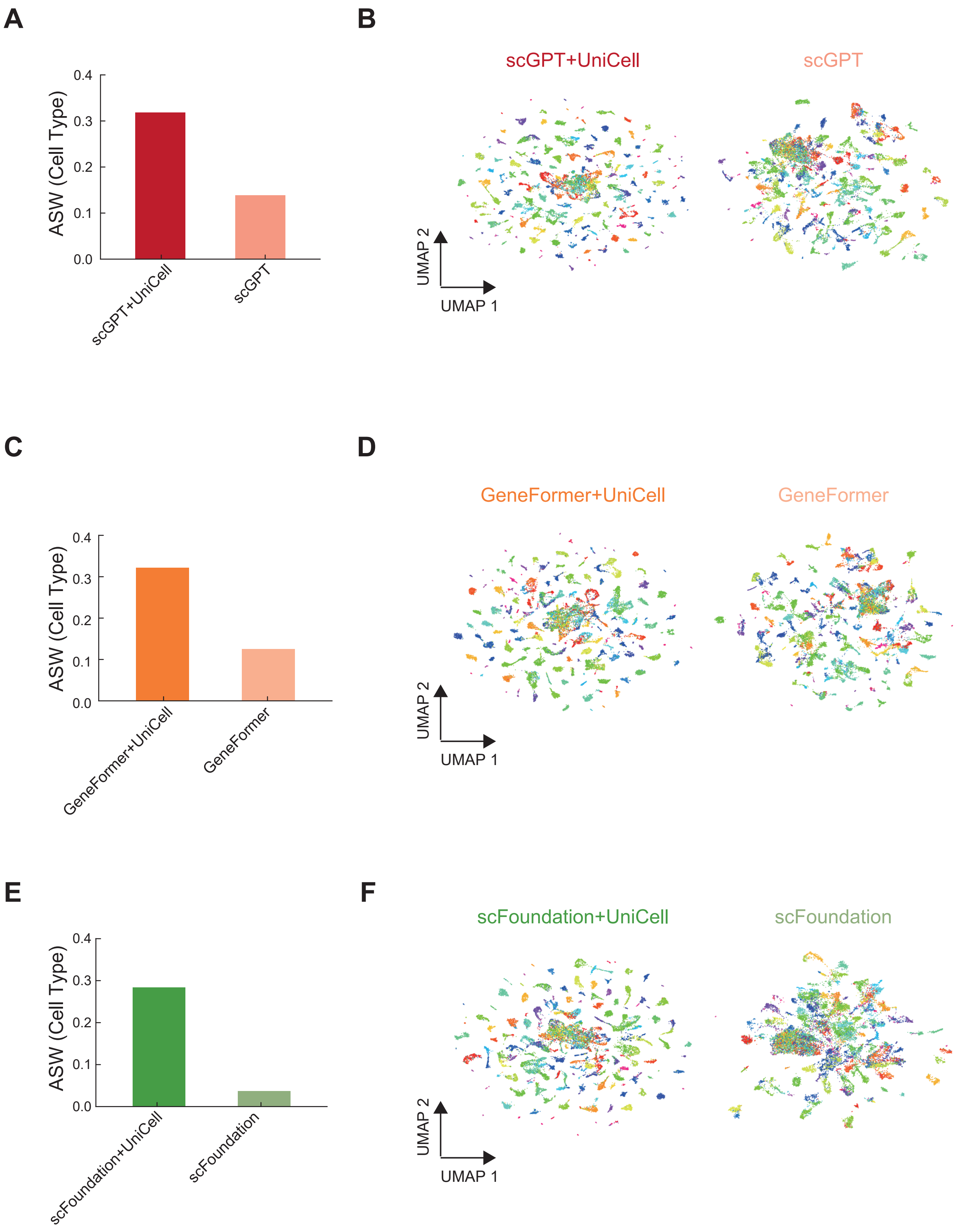

### Supplemental Figure 20

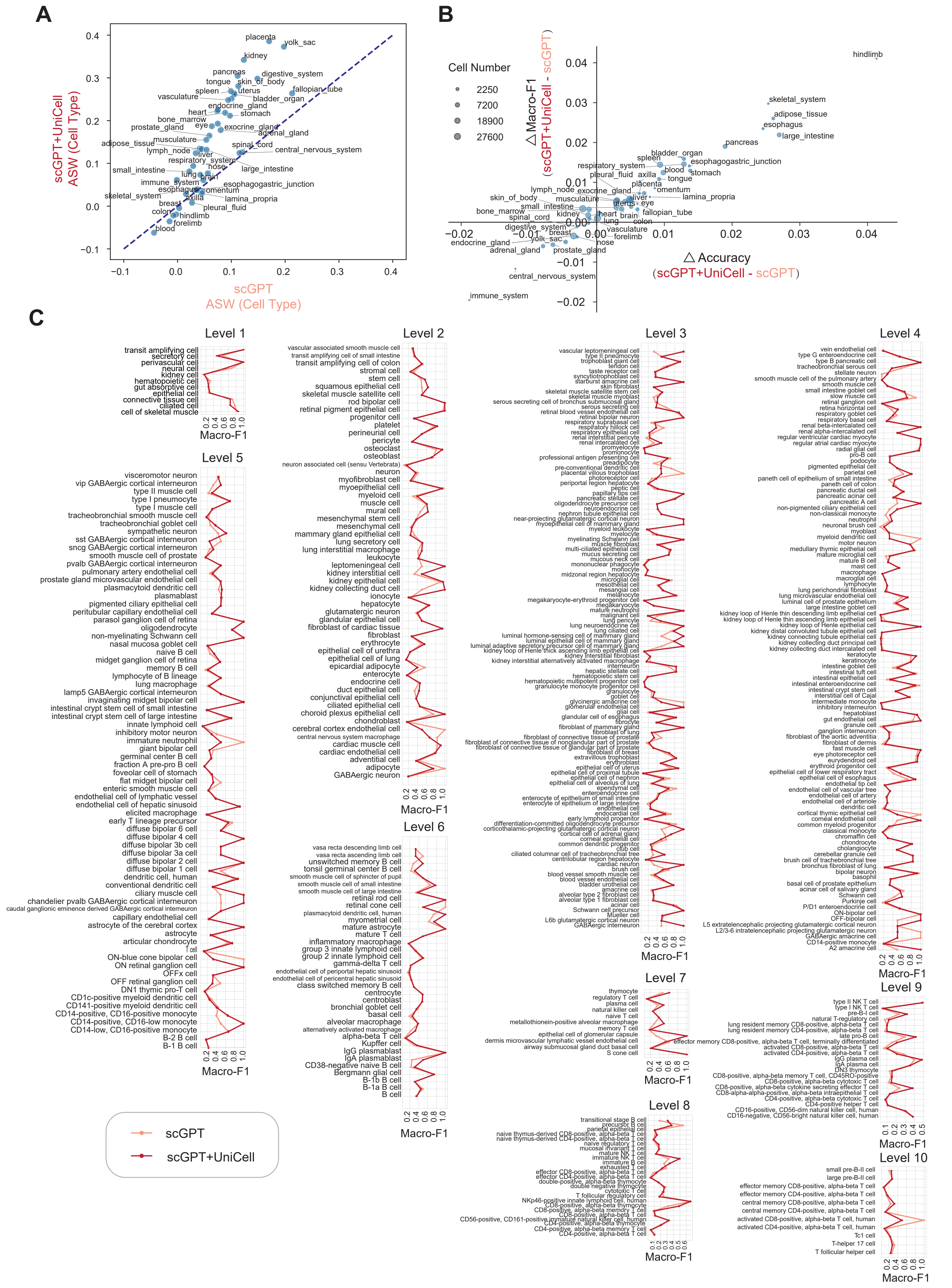
